# BATSY4-PRO: An Open-Source Multichannel Ultrasound Recorder for Field Bioacoustics

**DOI:** 10.1101/2025.08.11.669530

**Authors:** Ravi Umadi

## Abstract

**Background:** A variety of behavioural studies of echolocation and acoustic communication in free-flying bats depend on multichannel ultrasonic recordings. However, access to these methods remains geographically uneven: tropical and subtropical regions harbour much of the world’s bat diversity yet remain underrepresented in experimental behavioural bioacoustics. Conventional systems often depend on costly audio interfaces, laptops, and specialist expertise. Although instrumentation is only one contributor to this disparity, a compact, affordable, openly documented platform can reduce a practical barrier to field research, ecological monitoring, and conservation efforts.

**Results:** Here, I present Batsy4-Pro, an open-source four-channel recorder designed for behavioural and ecological field studies. Built around a Teensy 4.1 microcontroller and two WM8782 analogue-to-digital converters, it records synchronised 16-bit audio at 192 kHz directly to microSD storage. The recorder weighs under 150 g, operates from a 5 V supply, and, together with the four-microphone array, can be constructed for approximately 200 euros. It combines retrospective and buffer-prefilled forward recording with selectable real-time monitoring. While all four full-spectrum channels are recorded, users can monitor any individual microphone or a four-channel mix through tunable heterodyne conversion or direct passthrough. Field recordings showed consistently high signal-to-noise ratios comparable to recordings obtained with a commercially available multichannel soundcard and supported time-difference-of-arrival localisation of bat flight trajectories. Because the intended applications involve moving animals, I used grid-based Monte Carlo simulations to define the usable localisation field of the deployed array and evaluate how simulated source velocity affects localisation performance. The simulations identified a geometry-dependent near-array region of useful performance, followed by a sharp decline in accuracy with increasing range, and showed that localisation errors increased with simulated source velocity.

**Conclusions:** Batsy4-Proestablishes an open, reproducible workflow from synchronised field recording and real-time monitoring to three-dimensional source localisation. Its configurable firmware logic and extensibility allow the same compact system to be adapted to distinct behavioural questions, species, and habitats without dependence on laboratory-bound or proprietary multichannel platforms. Wider adoption can expand the geographic and taxonomic scope of experimental bat bioacoustics, strengthen local capacity for ecological monitoring, and enable comparative investigation of echolocation behaviour under natural conditions.

## 1 INTRODUCTION

Scientific progress often hinges on the availability of cost-effective, portable, and easily customisable tools. In recent years, the growing ecosystem of open-source electronics has played a pivotal role in enabling non-commercial development of field instruments, acting as a bridge between engineering innovation and ecological research (Oellermann et al., 2022; Pearce, 2020). These community-supported platforms empower scientists to prototype, iterate, and deploy experimental hardware tailored to specific scientific questions, especially in resource-constrained or remote settings (Ackermann, 2009; GOSH, 2017; Pearce, 2012, 2014; Powell, 2015).

Advances in embedded computing, particularly in high-performance microcontrollers, have opened new frontiers for field-based data acquisition. The domain of bioacoustics, especially studies involving ultrasonic signals in bats and insects, stands to benefit significantly from these developments. Despite the availability of compact recorders such as Solo (Whytock and Christie, 2017) and AudioMoth (Hill et al., 2019), and other open-source projects such as Raspberry Pi Bat Projekt (NABU e.V., 2016), most remain limited by proprietary hardware, closed firmware, or single-channel architectures that constrain their adaptability for advanced experimental designs requiring spatial localisation and sound-source tracking. Readily customisable, fully open-source multichannel ultrasonic recording systems with integrated real-time monitoring capabilities remain notably absent from the available toolkit (see Table 1). Recent work on inverse array design further indicates that task-specific placement of a small number of microphones can provide a practical alternative to indiscriminately increasing channel count, particularly when array aperture and portability are constrained (Umadi, 2026f).

**Table 1.**
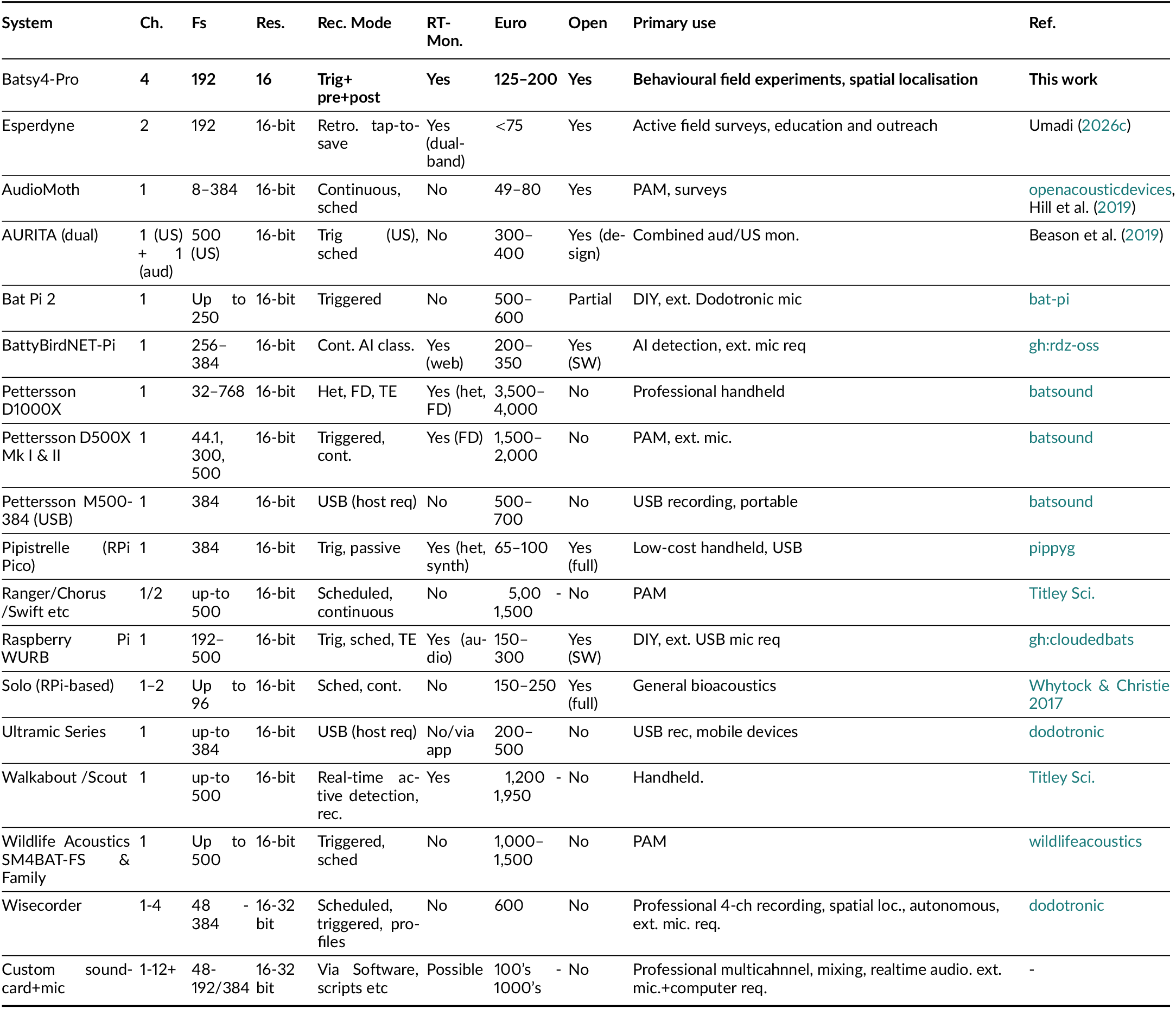
Comparison of Ultrasonic Recording Systems for Bat Bioacoustics Research. Batsy4-Pro offers a low-cost, fully open-source combination of: (1) true multichannel (4-channel) synchronous recording at 192 kHz, supporting spatial localisation; (2) integrated selectable real-time monitoring for immediate acoustic feedback during field deployment; (3) dual recording-mode architecture with retrospective and prospective capabilities for capturing spontaneous behavioural events; (4) modular, extensible firmware and hardware design; and (5) field-demonstrated acquisition performance comparable to commercial systems. Unlike passive monitoring devices designed for autonomous ecological surveys, the system is specifically engineered for active behavioural experiments requiring realtime decision-making, spatial resolution, and flexible triggering. This fills a critical gap in accessible tools for experimental bioacoustics research in resource-limited settings. *The costs are approximations based on publicly available information at the time of writing. The set of features is subject to manufacturer updates*.

Behavioural field experiments on echolocating bats face particular bottlenecks due to the lack of accessible, flexible, and field-ready multichannel recording systems. Traditional laboratory setups typically require professional audio interfaces, external preamplifiers, laptop computers, and stable AC power—configurations that are cumbersome to deploy, expensive to replicate, and require substantial technical expertise to operate reliably under field conditions. Ultrasonic microphones with flat frequency responses beyond 100 kHz add further cost. The cumulative cost of conventional multichannel field systems can readily reach several thousand euros when commercial recorders are combined with multiple localisation microphones, a calibrated measurement chain, and acoustic calibration equipment, as illustrated by a recent four-channel study of free-flying pipistrelles (Frommolt et al., 2026). These technical and economic demands can limit participation in experimental bioacoustics, particularly in tropical and subtropical regions where bat diversity is greatest but behavioural research remains underrepresented (Van Helden, 2012). Of the more than 1300 known species of echolocating bats, fewer than 30 have been studied in experimental behavioural contexts, reflecting both the technical demands and economic constraints of the field. Beyond its immediate utility for behavioural field studies, Batsy4-Pro is well aligned with emerging needs in computational bioacoustics, where large, well-annotated, high-quality acoustic datasets underpin advances in automated detection, classification, and behavioural inference (Mac Aodha et al., 2022; Stowell, 2022).

To address these limitations, I present Batsy4-Pro, an open-source, microcontroller-based four-channel ultrasonic recorder designed explicitly for behavioural field experiments with echolocating animals. Built on the Teensy 4.1 platform (Stoffregen and Coon, 2020), the system integrates all essential components—analogue-to-digital conversion, computational processing, data storage, and selectable real-time monitoring—into a compact device weighing less than 150 grams and operating from a standard USB power bank. The system records synchronised 16-bit audio at 192 kHz per channel using dual WM8782 ADCs, with data saved as timestamped WAV files to an onboard microSD card. External memory-based ring buffering enables dual-mode event-triggered recording: retrospective capture of the preceding 5 seconds and forward recording with prefill, which preserves behavioural context while continuing to record ongoing activity (terminology in Table 2).

**Table 2.** Abbreviations, Technical Terminology, and Audio Library Elements.

| Term | Description |
| --- | --- |
| **Hardware Components** |  |
| ADC | Analogue-to-Digital Converter. Converts analogue audio signals from microphones into digital data. The WM8782 is a 24-bit stereo ADC operating at up to 192 kHz. |
| OSR | Oversampling Ratio. The factor by which the ADC's internal sigma-delta modulator samples faster than the target output rate. The WM8782 uses 32× OSR at 192 kHz, meaning it samples internally at 6.144 MHz. |
| Sigma-Delta ADC | Type of ADC that uses oversampling and noise shaping to achieve high resolution. Converts analogue signals to a high-speed 1-bit stream, then decimates and filters to produce multi-bit output at the target sample rate. |
| DAC | Digital-to-Analogue Converter. Converts digital audio data back to analogue signals. The PCM5102A DAC is used for real-time monitoring output to headphones. |
| MEMS | Micro-Electro-Mechanical Systems. Miniature microphones (e.g., SPU0410LR5H-QB). |
| MCU | Microcontroller Unit. The Teensy 4.1 (ARM Cortex-M7, 600 MHz) serves as the central processing unit. |
| PSRAM | Pseudo-Static Random Access Memory. External memory used for the ring buffer, accessed via EXTMEM directive in C++. |
| I<sup>2</sup>S | Inter-IC Sound. Serial bus interface for digital audio transmission. Two I<sup>2</sup>S buses (I<sup>2</sup>S1, I<sup>2</sup>S2) connect the ADCs to the Teensy. |
| Slave Mode | ADC operation mode, where clock signals (BCLK, LRCLK) are provided by the master device (Teensy). The WM8782 operates in slave mode, with the M/S pin tied to logic low (0). |
| I<sup>2</sup>C | Inter-Integrated Circuit. Serial communication protocol used for the OLED display. |
| **Signal Processing** |  |
| DDS | Direct Digital Synthesis. Method for generating waveforms digitally using a phase accumulator, employed for sine wave carrier generation. |
| SNR | Signal-to-Noise Ratio. Measure of signal quality expressed in decibels (dB). |
| THD | Total Harmonic Distortion. Measure of nonlinearity in audio systems, expressed as a percentage or in decibels. Quantifies the ratio of unwanted harmonic frequencies to the fundamental frequency. |
| PCM | Pulse-Code Modulation; a digital representation of an analogue audio signal in which the signal amplitude is sampled at fixed time intervals and quantised into discrete numeric values. Used for lossless audio storage formats such as WAV files. |
| Heterodyne | Frequency mixing technique that multiplies an ultrasonic signal with a carrier sine wave to produce audible frequencies. |
| Phase Accumulator | Digital oscillator that generates a sine wave by incrementing a phase variable per sample. |
| TDoA | Time-Difference of Arrival. Method for spatial localisation using arrival time differences between microphones. |
| **Teensy Audio Library Elements** |  |
| AudioInputI2S | Audio input object for the first I<sup>2</sup>S interface (I<sup>2</sup>S1). Receives stereo data from ADC 1 (channels L1, R1). |
| AudioInputI2S2 | Audio input object for the second I<sup>2</sup>S interface (I<sup>2</sup>S2). Receives stereo data from ADC 2 (channels L2, R2). |
| AudioRecordQueue | Buffer object that stores incoming audio blocks (128 samples each) for processing. Four instances used: queueL1, queueR1, queueL2, queueR2. |
| AudioPlayQueue | Buffer object for audio playback. Used to send monitoring samples to the DAC output. |
| AudioOutputI2S | Audio output object that transmits digital audio to the DAC via I<sup>2</sup>S. |
| AudioConnection | Defines signal routing between audio objects. Example: patchCord1(i2sInput1, 0, queueL1, 0) connects I<sup>2</sup>S1 left channel to queue. |
| AudioMemory() | Allocates memory blocks for audio processing. AudioMemory(120) reserves 120 blocks for buffering. |
| readBuffer(), getBuffer(), playBuffer() | Audio queue buffer management methods used to retrieve input sample blocks, allocate output buffers, and submit filled buffers for digital-to-analogue conversion during real-time audio processing. |
| available() | Method to check if a queue has data ready for reading or if an output buffer is available. |
| clear() / freeBuffer() | Methods to release processed audio blocks back to the memory pool after use. |
| **Recording Modes** |  |
| Retrospective (TAP) | Recording mode that saves the previous 5 seconds from the ring buffer when triggered by the tap button. |
| Forward with Prefill (HOLD) | Recording mode that first saves 5 seconds of buffered audio, then continues recording up to 10 seconds of new audio when the hold button is pressed. |
| **Clock Configuration** |  |
| PLL | Phase-Locked Loop. Hardware oscillator that generates precise clock signals for I<sup>2</sup>S interfaces. |
| SAI | Synchronous Audio Interface. Teensy 4.1 peripheral that implements I<sup>2</sup>S communication (SAI1 = I<sup>2</sup>S1, SAI2 = I<sup>2</sup>S2). |
| BCLK | Bit Clock. Clock signal that times individual bit transmissions in I<sup>2</sup>S protocol. |
| LRCLK | Left-Right Clock (Word Clock). Indicates whether left or right channel data is being transmitted. |

A key feature distinguishing Batsy4-Pro from passive acoustic monitoring devices is its integrated real-time audio monitoring system. It provides tunable heterodyne monitoring of ultrasonic signals or direct passthrough of audible-frequency signals. The output remains active during recording and provides immediate acoustic feedback through standard headphones, enabling manual triggering from observed call patterns without a separate detector or real-time spectrogram display.

Because Batsy4-Pro is intended for recording free-flying bats, characterising the deployed array requires consideration of source movement as well as microphone geometry. Localisation error propagates directly into velocity estimates derived from successive acoustic positions, while source motion can itself alter localisation performance. These coupled effects mean that position and velocity estimates should be interpreted only after localisation uncertainty has been characterised across the intended field of interest (Umadi, 2026f). This issue is exemplified by a study in which flight speed was calculated from successive acoustically localised source positions without separately quantifying the contribution of velocity-dependent localisation error (Jakobsen et al., 2025). I therefore applied the Widefield Acoustics Heuristic framework to determine the useful spatial field of the realised array and examine its sensitivity to source velocity (Umadi, 2025d).

I evaluated Batsy4-Pro through a controlled functional playback test and field deployment with free-flying bats in a natural foraging corridor. The field recordings consistently showed high signal-to-noise ratios (median 27.3 dB), comparable to those obtained with a professional-grade audio interface, and provided synchronised multichannel signals suitable for time-difference-of-arrival localisation. Simulations were used separately to characterise the expected spatial limits of the demonstrated array. The total component cost, including recorder and four-channel microphone array, is approximately 200 euros when sourced locally. All hardware designs, circuit schematics, firmware source code, and assembly documentation are released under permissive open-source licensing^1^ (Umadi, 2026b) that explicitly permits commercial production without royalty obligations, facilitating local manufacturing and institutional workshop projects.

Batsy4-Pro demonstrates that high-quality multichannel ultrasonic recording for experimental bioacoustics can be achieved using affordable, accessible, open-source hardware. By combining field-tested performance with modular extensibility and comprehensive documentation, this work aims to lower technical and economic barriers to behavioural research with echolocating animals.

## 2 METHODS

This section describes the design, implementation, and performance characterisation of the Batsy4-Pro system, combining hardware architecture, firmware design, and analytical methods used to evaluate its operation under field-relevant conditions. I first detail the recorder’s electronic design, signal acquisition pipeline, buffering strategy, and selectable real-time monitoring. I then describe the construction and deployment of the four-channel microphone array, followed by laboratory and field demonstrations. Finally, I outline the simulation analysis used to characterise spatial localisation uncertainty and the expected operational volume of the demonstrated array for behavioural bioacoustics experiments.

### 2.1 Design and Implementation

The following section outlines the hardware architecture, signal-processing workflow, and firmware structure of the Batsy4-Pro ultrasonic recorder – provided to document the system’s design and performance; familiarity with all technical aspects is not required for routine use or deployment. Definitions of technical terms, abbreviations, and the *Teensy Audio Library* components are summarised in Table 2.

Batsy4-Pro is a four-channel ultrasonic audio recorder built around the Teensy 4.1 microcontroller unit (MCU) (Stoffregen and Coon, 2020). It is designed for field deployment in behavioural studies of echolocating animals. The system records synchronised 16-bit audio at 192 kHz per channel and saves data as WAV files to an onboard microSD card. This sampling rate gives a theoretical Nyquist frequency of 96 kHz; the usable acoustic band-width below this limit is additionally determined by the microphone response and analogue filtering. Two tactile buttons enable user control: one triggers retrospective capture of the preceding 5 s from a ring buffer, while the other initiates a recording containing the preceding 5 s followed by up to 10 s of live audio. The firmware is written in C++ using the Arduino IDE environment and leverages the *PJRC Teensy Audio library* (Stoffregen and Coon, 2015); the complete implementation is available in the accompanying repository (Umadi, 2026a). A rotary encoder selects the monitoring mode, input, and output level. The carrier control provides heterodyne monitoring from 10–85 kHz or direct passthrough, while the input control selects Channels 1–4 or their combined mix (Figure 1a).

**Figure 1.**
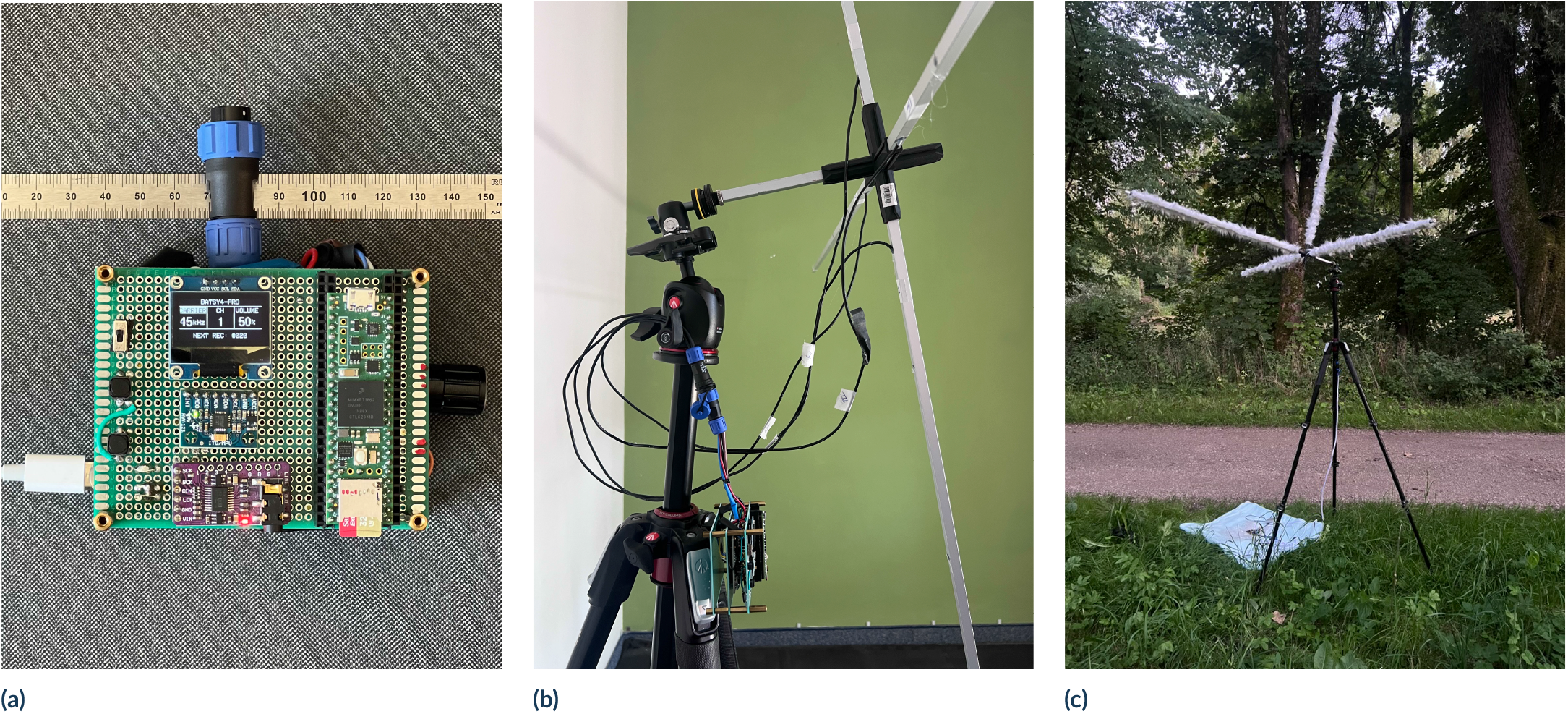
The Batsy4-Pro recorder and field array. (a) Recorder prototype assembled on perforated PCB, with components soldered in two stacked layers. External 5 V power is supplied through a USB-C socket, and a detachable 6-pin connector links the recorder to the array microphones. Four-channel recordings are stored on the onboard microSD card, while selectable direct-passthrough and 10–85 kHz heterodyne monitoring can be heard through conventional headphones. Component functions and signal flow are detailed in Figure 2. (b) Rear view of the array hub showing the 6-way connector, internal wiring, detachable arm sockets, and an aluminium mounting rod with a 1/4” tripod thread attached to a ball head. (c) Fully assembled cross-type microphone array mounted on a tripod at the field site. The four 1 m arms occupy hub sockets along three dimensions; each microphone port is positioned 1 cm inboard from the distal arm tip, giving a hub-to-microphone distance of 0.99 m. The detachable arms can be inserted into different hub sockets, as shown by the configurations in (b) and (c).

#### 2.1.1 Components

The core hardware components include:

- Teensy 4.1 microcontroller unit (600 MHz ARM Cortex-M7, 16-bit audio support)
- Two APS6404L-3SQR PSRAM 8MB (Soldered on Teensy 4.1), 16MB allows extending forward record durations to 20 s.
- Two WM8782 stereo ADCs interfaced via I^2^S1 and I^2^S2
- PCM5102A-based DAC Module with a headphone jack
- Four Knowles SPU0410LR5H-QB analogue omnidirectional MEMS microphones (direct differential input to ADCs)
- Adafruit SSD1306 128 × 64 pixel OLED display (I^2^C interface)
- FAT32-formatted microSD card inserted into the builtin Teensy slot for data storage
- Two tactile push buttons: For retrospective and prospective recording.
- A rotary encoder with switch function: selecting the carrier, input-channel, or output-level control and adjusting heterodyne frequency or passthrough (`PT`) mode, Channels 1–4 or their mix (`MX`), and monitoring volume.

A range of MEMS and conventional analogue microphones can meet different recording requirements, provided that the selected sensor is compatible with the ADC input and system power supply. Application-specific performance evaluation is encouraged when substituting the microphone.

Power is supplied via USB from a portable battery pack. Refer to Supplementary Table T2 for the recorder bill of materials and Supplementary Table T1 for component-to-component and component-to-Teensy connections.

#### 2.1.2 Circuitry

The system architecture and signal flow are presented in figure 2. Each ADC interfaces with the Teensy over dedicated I^2^S lines. Audio connections are managed via ‘AudioInputI2S’ and ‘AudioInputI2S2’ classes. A pair of ‘AudioRecordQueue’ objects buffers each input channel. Audio buffers are written as interleaved 16-bit samples into a WAV file stream on the SD card.

**Figure 2.**
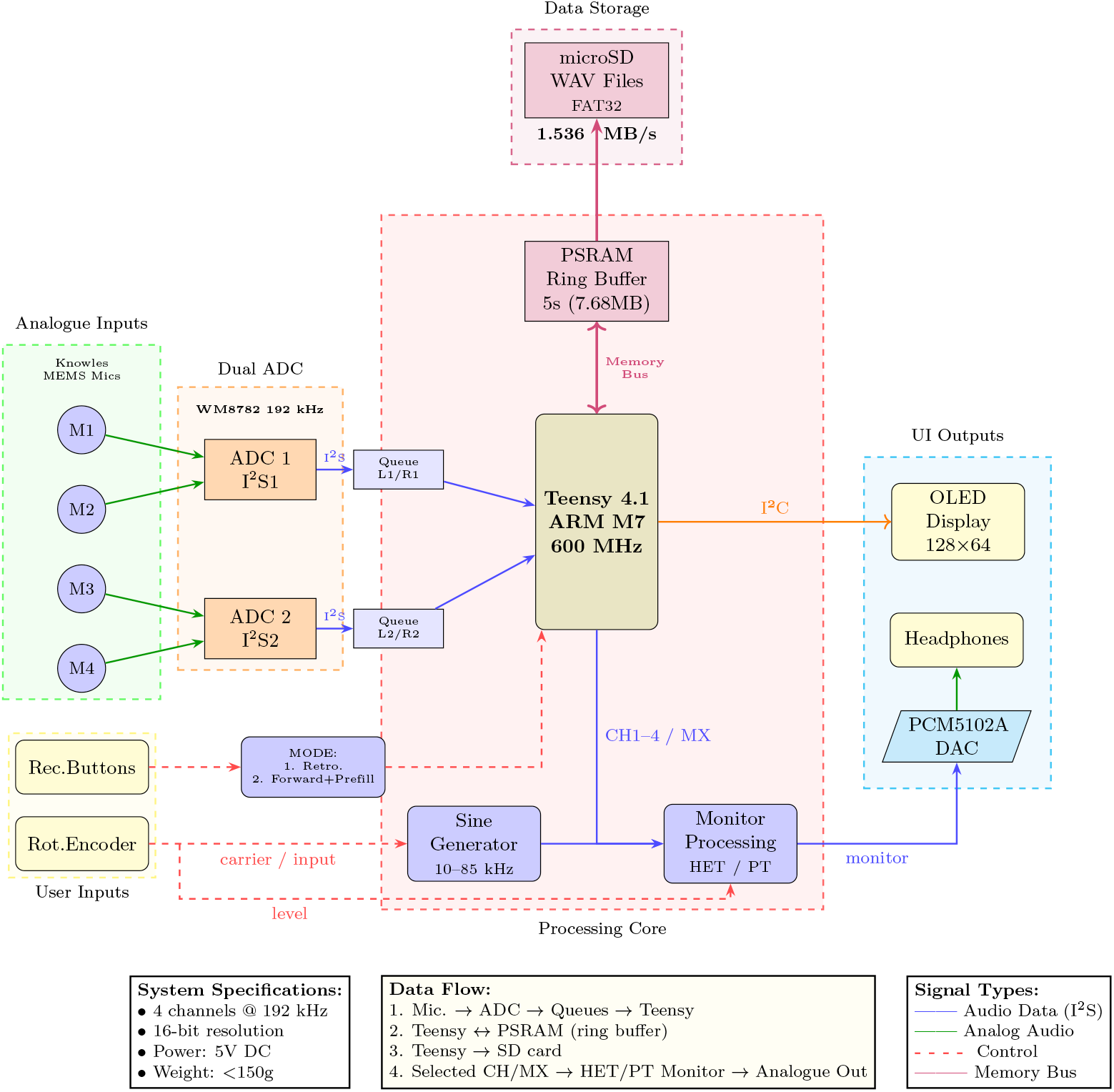
System architecture and data flow of Batsy4-Pro. Two WM8782 stereo ADCs digitise four analogue microphones (M1–M4) at 192 kHz and stream via dual I^2^S interfaces into a Teensy 4.1 microcontroller (ARM Cortex-M7, 600 MHz). Audio samples are buffered in channel-specific queues and written to an external PSRAM ring buffer, enabling both retrospective and forward-with-prefill recording modes. Upon user trigger, buffered data are interleaved and written as time-stamped, four-channel WAV files to a FAT32-formatted microSD card. In parallel, an operator-selected input (Channels 1–4 or their combined mix, `MX`) is routed either through the heterodyne mixer or directly to the DAC in passthrough (`PT`) mode for real-time headphone monitoring; Channel 1 with a 45 kHz carrier is selected by default. System state, monitoring settings, and recording status are displayed on an OLED screen.

### 2.2 I^2^S ADC Configuration

The system uses two WM8782 ADC boards, each capable of digitising stereo analogue signals at up to 24-bit / 192 kHz. These are configured to operate in *slave* mode, streaming data via I^2^S to the Teensy 4.1’s `I2S1` and `I2S2` ports simultaneously, enabling synchronised four-channel audio recording. Each ADC contributes two 16-bit stereo channels per frame, with the PJRC Audio library managing the dual I^2^S streams.

The WM8782 is a 16/20/24-bit sigma-delta stereo ADC designed for high-fidelity audio capture. It supports sample rates from 8 kHz to 192 kHz and offers digital audio output in I^2^S, left-justified, or right-justified format. Its performance includes a signal-to-noise ratio (SNR) of up to 100 dB and total harmonic distortion (THD) better than −93 dB at 48 kHz (*WM8782 24-Bit 192 kHz Stereo ADC Product Data Sheet* 2020).

Each ADC operates in *slave mode*, meaning that bit clock (BCLK) and word clock (LRCLK) are provided by the Teensy. The WM8782 automatically detects the sampling rate by locking onto these clock signals, with a built-in mechanism to determine the oversampling ratio (OSR) via the `FSAMPEN` pin. To match the 192 kHz operation of the system, `FSAMPEN` is set to high-impedance (Z), resulting in 32× oversampling (*WM8782 24-Bit 192 kHz Stereo ADC Product Data Sheet* 2020).

#### 2.2.1 ADC Configuration Pins

The WM8782 is fully hardware-configured via digital input pins as follows:

- `FORMAT` (Pin 9): set to high-impedance (Z) for I^2^S format
- `IWL` (Pin 7): set to low-impedance (Z) for 16-bit word length. Although the WM8782 supports 24-bit resolution, the current implementation uses 16-bit to maintain compatibility with the *Teensy Audio Library*.
- `M/S` (Pin 20): tied to logic low (0) for slave mode
- `FSAMPEN` (Pin 8): set to high-impedance (Z) to enable 192 kHz sampling (32× OSR)

This configuration results in each WM8782 transmitting 16-bit I^2^S audio data synchronised to the Teensy’s clocks, ensuring sample-accurate alignment across all four channels.

### 2.3 I^2^S Sampling Rate Configuration

Precise sampling rate configuration is critical in multichannel ultrasonic systems to ensure temporal synchrony across input channels and to preserve waveform fidelity at high frequencies. To achieve this, both the software sampling definitions and the Teensy’s hardware phase-locked loop (PLL) clocks are explicitly configured to deliver a stable sampling rate of 192 kHz per channel across both I^2^S interfaces.

#### 2.3.1 Audio Library Sampling Rate

To ensure the Teensy Audio library internally aligns with the desired high-resolution sampling, the sampling rate is explicitly defined using:

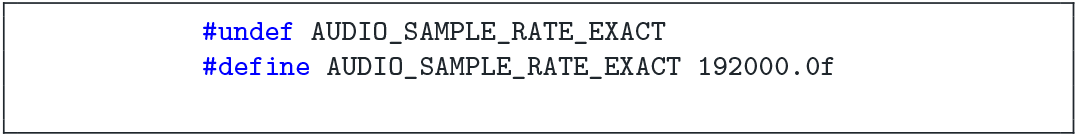

This directive ensures that all internal buffers, sample timing, and I^2^S input and output objects are configured for a 192 kHz sample rate before any audio object initialisation.

#### 2.3.2 Master Clock Settings

The following clock configuration is a critical enabling step for high-frequency, multi-channel ultrasonic recording on the Teensy 4.1. To achieve a stable sampling rate of 192 kHz per channel, a custom clock configuration is implemented through the `setI2SFreqBoth()` function (Supplementary Section A). This function configures the internal phase-locked loop (PLL) on the Teensy 4.1 to generate the precise master clock required for high-rate I^2^S communication on both SAI1 and SAI2 interfaces (*i.MX RT1060 Processor Reference Manual* 2021).

The function first calculates the necessary PLL multiplier and divider settings for a given sampling frequency. The core calculation derives from the formula:

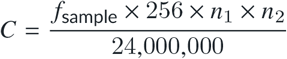

Where *f*_sample_ is the target sampling frequency (192 kHz), *n*_1_ is the SAI prescaler (set to 4), and *n*_2_ is the PLL divider computed to approximate the required master clock. The fractional components of *C* are then passed to the `set_audioClock()` function, which configures the audio clock source with the appropriate numerator and denominator.

After configuring the PLL, the function sets the I^2^S peripheral clock dividers (`CCM_CS1CDR` and `CCM_CS2CDR`) for both I^2^S1 and I^2^S2. This ensures that both stereo input channels operate synchronously at the same sample rate. This configuration guarantees a sampling rate of exactly 192000 Hz for all four input channels (two per I^2^S interface), maintaining synchrony and timing accuracy required for multichannel ultrasonic recordings (see Chapters 14 and 38 of the i.MX RT1060 Crossover Processor reference manual for a detailed system description on Clock Control Module and SAI (*i.MX RT1060 Processor Reference Manual* 2021)).

### 2.4 Memory Buffer Architecture and Recording Modes

The firmware implements two complementary data capture modes that are directly referenced in the user interface and code as **TAP** and **HOLD** modes. These correspond to (i) a *retrospective buffer-triggered capture* (TAP), which saves a fixed-duration (default 5 s) window of recent audio from a continuously running ring buffer, and (ii) a *forward-triggered capture with buffer prefill* (HOLD), which appends ongoing audio recording to a preceding segment of buffered history. Together, these modes enable both rapid reaction to transient events and extended recording while preserving temporal context. The flowchart in figure 3 presents the implementation logic.

**Figure 3.**
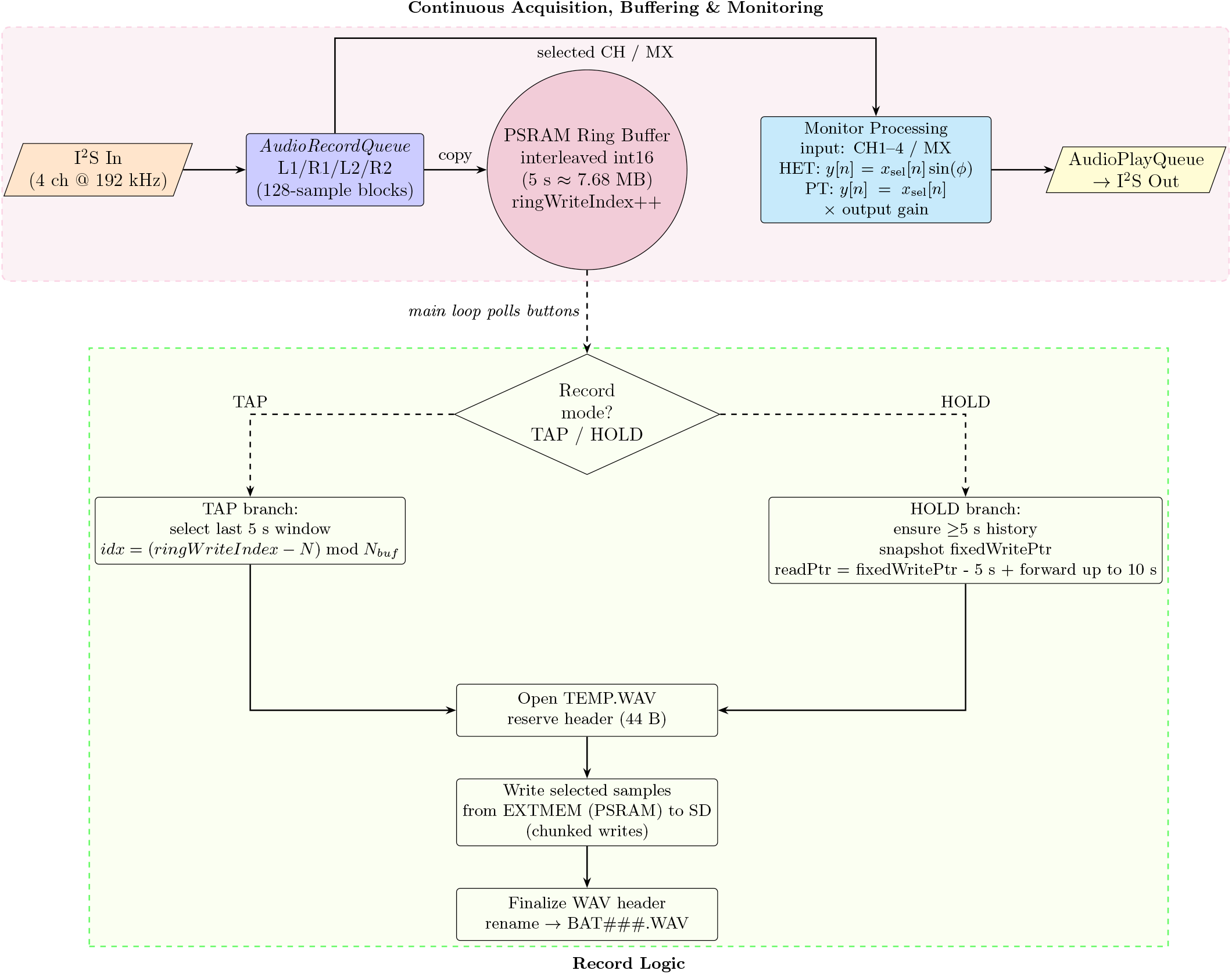
Ring-buffer recording logic. Four synchronised 192 kHz channels are continuously acquired in 128-sample blocks via `AudioRecordQueue` objects and interleaved into a 5 s PSRAM (`EXTMEM`) ring buffer, while a user-selected channel or four-channel mix is simultaneously routed through heterodyne or passthrough processing for real-time monitoring. The main loop polls the user inputs to select the capture mode: **TAP** triggers *retrospective* capture by computing the start index for the last 5 s window *idx*, and copying that segment from PSRAM to a temporary WAV file using chunked writes; **HOLD** triggers *buffer-prefilled forward* capture by first ensuring 5 s history is available, snapshotting `fixedWritePtr`, writing ≥5 s of pre-buffer, then appending up to 10 s of forward audio before finalising. In both modes, audio is written to `/TEMP.WAV` with a reserved 44-byte header region, then the header is populated, and the file is renamed to a unique identifier.

The 5 s retrospective duration was selected as an engineering compromise between retaining pre-trigger behavioural context and the memory required for four synchronised channels. At 16-bit resolution and a sampling rate of 192 kHz, this window occupies 7.68 MB of PSRAM. It also encompasses the brief terminal approach and interception sequences analysed in representative bat-prey studies, which focus on the final few seconds of rapidly un-folding behaviour (Jakobsen et al., 2025; Xia et al., 2025).

When a longer sequence is required, the buffered forward-recording mode retains the same 5 s pre-trigger history and appends up to 10 s of live audio. This capture strategy has also proved sufficient for targeted field recordings of water-foraging bats (Umadi, 2026d). The retrospective duration remains configurable in the firmware and can be extended when lower sampling rates or fewer channels reduce the memory requirement.

#### 2.4.1 Ring Buffer

Digitised samples arriving from the microphones through the ADC signal chain are collected in 128-sample blocks for each channel using `AudioRecordQueue` objects. The channel blocks are interleaved and written sequentially to a rolling ring buffer in external PSRAM (`EXTMEM`). Figure 3 summarises this continuous acquisition, buffering, and capture workflow.

Under the recording configuration described above, storing each sample as 16-bit PCM produces a sustained data rate of

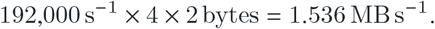

The resulting 5 s history occupies 7.68 MB of PSRAM. A circular write index advances continuously and overwrites the oldest samples once the buffer is full, while a cumulative sample counter ensures that a complete pre-trigger history is available before capture can begin. Further implementation details are provided in Supplementary Section B.

#### 2.4.2 TAP Mode: Retrospective Buffer Capture

In **TAP mode**, a user tap triggers immediate saving of the most recent 5 seconds of audio from the ring buffer. Upon activation, the firmware computes a read pointer offset relative to the current write (to buffer) index and sequentially copies the buffered samples into a temporary WAV file (`/TEMP.WAV`). Data are written in small, non-blocking chunks (128–512 samples), interleaved with ongoing audio acquisition and processing, so that live recording and monitoring continue uninterrupted.

Once the buffered window has been written, a standard 44-byte WAV header is inserted, and the file is finalised and renamed using an incrementing identifier (e.g., `/BAT001.WAV`). TAP mode, therefore, functions as a rolling “look-back” recorder, allowing capture of brief or unexpected acoustic events that occurred prior to user intervention. Since the buffer is not deleted after recording, subsequent TAP triggers within 5 s of the last recording include the newest data, with overlaps from the previous recording.

#### 2.4.3 HOLD Mode: Forward Recording with Buffer Prefill

In **HOLD mode**, recording begins with a retrospective prefill followed by forward recording. When HOLD is activated, the firmware proceeds once the ring buffer contains 5 seconds of audio history; if activated earlier, acquisition continues until the complete pre-trigger window is available. A snapshot of the current write index is then taken, establishing a fixed reference point for the pre-trigger segment.

The last 5 seconds of buffered audio are written to a new temporary WAV file, after which live audio is appended in real time for up to 10 seconds (user-configurable). During this forward phase, the firmware periodically compares the ring buffer write pointer and read pointer to determine how many new samples are available for flushing to disk, ensuring continuous recording without overruns or data loss. Recording terminates automatically upon timeout, after which any remaining buffered samples are written, and the WAV file is finalised.

#### 2.4.4 Operational Characteristics

Both TAP and HOLD modes operate within a single co-operative processing loop without interrupts or blocking calls. The ring buffer continues to update at all times, independent of recording state, and user input determines how buffered data are extracted and committed to storage.

This architecture provides reliable, low-latency, retrospective and prospective recording and continuous operation, making it well-suited for field deployments where acoustic events are brief and spontaneous.

### 2.5 Integrated Real-Time Monitoring

The system incorporates continuous real-time monitoring (continuous loop in Figure 3) for auditory assessment during field recordings.

#### 2.5.1 Monitoring Implementation

The monitor uses the same 192 kHz digitised stream as the full-spectrum recording. Channel 1 is selected by default at power-up, and the rotary encoder allows the operator to select Channels 1–4 or `MX`, which sums all four inputs before output processing. The carrier control selects heterodyne monitoring between 10–85 kHz or `PT`, which bypasses frequency conversion and routes the selected input directly to the DAC. Pressing the encoder cycles among carrier, monitored-input, and output-level controls; rotation changes the highlighted parameter shown on the OLED display. These settings affect only the monitoring output, while all four channels continue to be recorded concurrently as full-spectrum data. The default carrier is 45 kHz.

Carrier generation and sample-wise mixing use the phase-accumulator approach described for the ESPERDYNE dual-band heterodyne monitor (Umadi, 2026c), following established digital heterodyning principles (Leis, 2011; Tan and Jiang, 2013; Tierney et al., 1971). The resulting difference-frequency component maps ultrasonic energy into the audible range; for example, a 62 kHz component mixed with a 60 kHz carrier yields a 2 kHz tone.

No explicit digital low-pass filtering is applied after mixing. As in conventional heterodyne bat detectors, the high-frequency image band is naturally attenuated by the reconstruction filter of the PCM5102A digital-to-analogue converter (Texas Instruments, 2015) and by the downstream analogue playback chain. Because the heterodyne signal is used exclusively for real-time auditory monitoring and not for quantitative analysis, this minimal filtering approach is sufficient and avoids additional computational overhead.

The monitoring signal is constrained to 16-bit signed integers and streamed via an `AudioPlayQueue` to the `AudioOutputI2S` interface for playback through the DAC. Output amplitude is scaled by a user-adjustable gain parameter, effectively providing a digital volume control. All four input channels are simultaneously recorded in parallel as raw, full-bandwidth ultrasonic data to the ring buffer and SD card, ensuring that monitoring does not alter or interfere with the stored recordings. Full implementation details are provided in Supplementary Section C. Representative laboratory and field recordings are shown in Figure 4.

**Figure 4.**
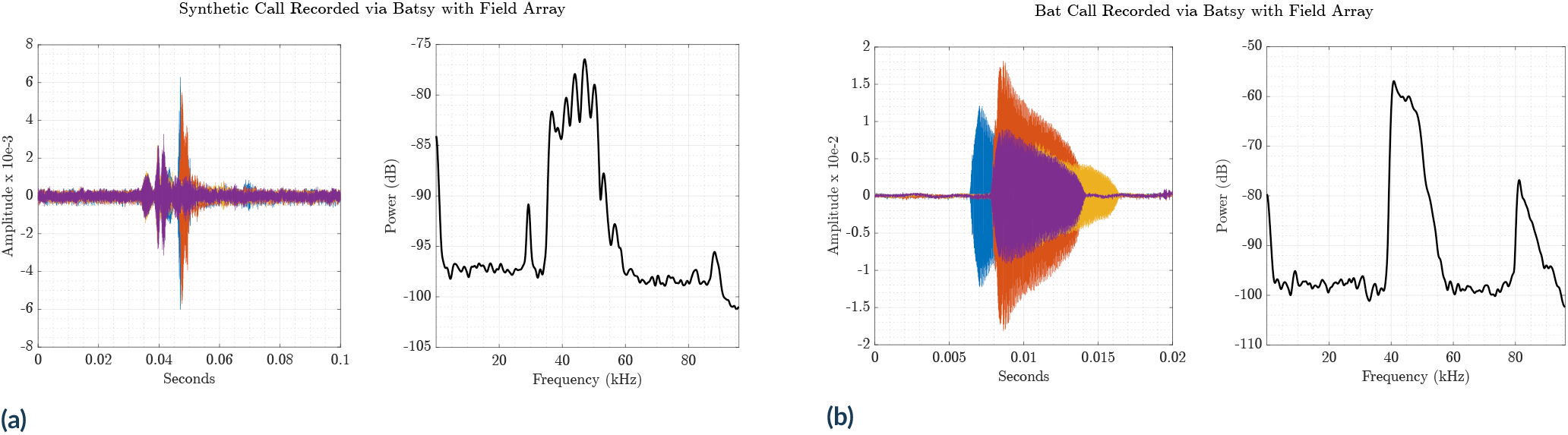
**Left (a):** A computer-generated quadratic FM chirp recorded indoors without acoustic insulation on either the array or the room, resulting in spectral interference from multipath reflections. The source was positioned approximately 2 m from the microphones. **Right (b):** A representative bat echolocation call recorded with the Batsy4-Pro system using the four-channel field array. The time-domain waveforms show all four channels, and the corresponding power spectrum was calculated from Channel 1. The spectrum reveals a prominent third harmonic of the echolocation call.

### 2.6 Array Assembly and Deployment

The microphone array consists of four Knowles SPU0410LR5H-QB analogue MEMS microphones with omnidirectional sensitivity, mounted on hollow aluminium arms arranged in a three-dimensional cross configuration (Figure 1c). Each arm is a 1 m-long, 1 cm^2^ square-section aluminium rod inserted into a central six-way connector hub (commercially available; see Supplementary Table T3). The microphone port is mounted 1 cm from the distal tip of each arm, so each microphone lies 0.99 m from the hub centre. The realised arrangement has one upper arm, one rearward arm, and two opposing lateral arms: adjacent non-collinear arms are separated by 90°, while the two lateral arms are separated by 180°. With Microphone 1 as the coordinate origin, the positions used for localisation and simulation were M1 (0, 0, 0), M2 (0, −0.99, −0.99), M3 (−0.99, 0, −0.99), and M4 0.99, 0, 0.99 m; equivalently, the hub lies at 0, 0, 0.99 m in this reference system. The microphones are wired internally along each arm, and all connections terminate at a single 6-pin connector at the hub, enabling fast, secure attachment to the recorder.

The central hub features a rear-mounted 15 cm solid aluminium rod (1 cm^2^ cross-section) with a custom-machined 1/4-inch tripod thread. This allows attachment to a standard ball-head adapter with a hot shoe plate mounted on a conventional tripod (Figure 1b). The assembled array, including the connector and mounting rod, weighs approximately 800 grams and is easily transportable.

During deployment, the arms are pulled out of the hub and inserted as needed, enabling rapid assembly and disassembly in field conditions. Each arm is fitted with tailored sleeves made of acoustic insulation material made of polyester fur.

The array geometry enables spatial localisation of ultrasonic calls when combined with synchronised four-channel recording. The precise configuration and rigid construction ensure repeatable microphone placement and stable performance during field sessions.

### 2.7 Laboratory and Field Demonstration

A laboratory functional test used synthetic FM chirps played through an RME Babyface (Audio AG, Haimhausen, Germany) audio interface, amplified via a single-channel power amplifier, and reproduced with a Scan-Speak R2004 tweeter (Scan-Speak A/S, Denmark). Playback level was not calibrated because the test assessed general recording functionality and relative signal-to-noise characteristics, rather than calibrating microphone sensitivity or reporting absolute sound-pressure levels. As expected in the untreated room, the recordings exhibited a complex pattern of reflections from surrounding objects. The test confirmed proper system operation before field deployment (Figure 4a).

To demonstrate a field use case and the functional deployment of the Batsy4-Pro system, a field experiment was conducted at a location known for frequent bat foraging activity. No special permit was necessary to record free-flying bats. The array was mounted on a tripod and deployed in a semi-open corridor clearing (Figure 1c).

Approximately 30 distinct foraging sequences characterised by rapid echolocation call bouts and frequency-modulated signals were recorded. Recordings were manually screened and sorted for high-quality multi-channel segments with high signal-to-noise ratios across all four microphones (Figure 4b) using the *Bat Reviewer* tool (Umadi, 2025b). 20 ms regions around selected calls were extracted via custom-written MATLAB scripts, adopted from the `Biosonar Responsivity Analysis Toolkit V1.0` (Umadi, 2025c).

### 2.8 SNR characterisation of field data

To quantify relative signal-to-noise performance under field conditions, extracted echolocation calls were analysed using a consistent, peak-based signal-to-noise metric (Figure 5). Noise was estimated from the quietest 2 ms window within the final portion of each clip, identified as the minimum root-mean-square (RMS) amplitude using a sliding-window search to minimise contamination by call energy.

**Figure 5.**
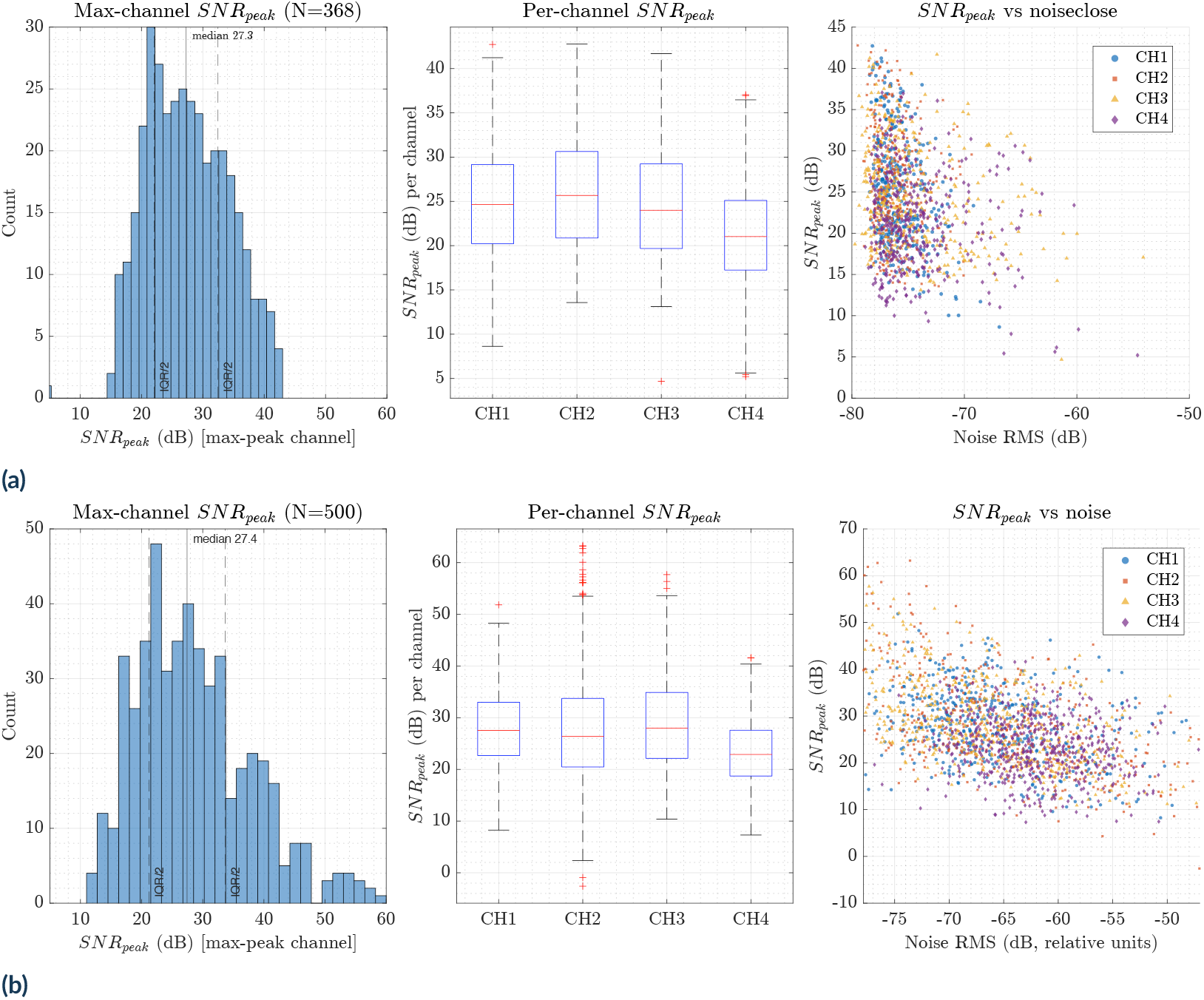
SNR Comparison against a professional audio interface. Peak-based signal-to-noise ratios (SNR_peak_) are shown for field recordings obtained with Batsy4-Pro (top). The bottom row shows a randomly selected subset of recordings obtained with a professional audio interface (Behringer U-PHORIA UMC404HD) from the field dataset reported in (Umadi and Firzlaff, 2025). For each system, panels show (A) the distribution of maximum-channel SNR_peak_ across calls, with dashed lines indicating the median and interquartile range, (B) per-channel SNR_peak_ distributions, and (C) the relationship between SNR_peak_ and noise root-mean-square (RMS) level for all calls and channels. Median maximum-channel SNR_peak_ values were nearly identical between systems (27.3 dB for Batsy4-Pro; 27.4 dB for the Behringer soundcard), and both datasets show similarly tight noise distributions.

For each channel, call amplitude was defined as the maximum absolute sample value outside the noise-search region. A peak-based signal-to-noise ratio (SNR_peak_) was computed as

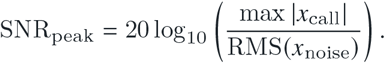

For each call, the channel exhibiting the highest peak amplitude was used to summarise maximum-channel SNR_peak_, while per-channel SNR distributions were analysed to assess channel-to-channel consistency.

### 2.9 Clustering and Localisation Methods

I implemented a combination of spectral clustering and time-difference-of-arrival (TDoA) based localisation to analyse the diversity and spatial structure of the recorded echolocation calls. The recorded call sequence, clustering results, and representative cluster members are summarised in Figure 6.

**Figure 6.**
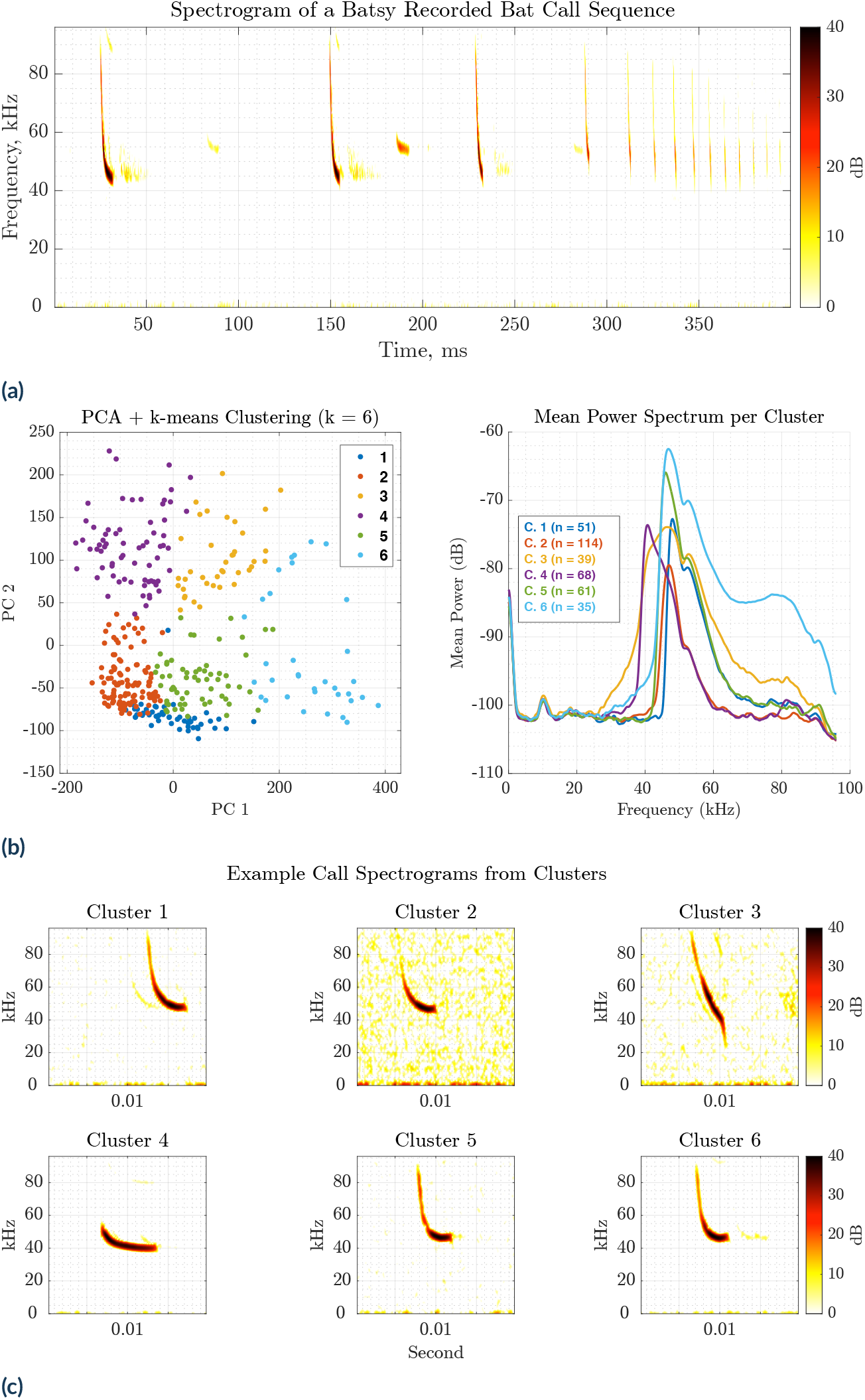
Field-data demonstration: (**a**) Spectrogram of a sequence of bat calls recorded on the Batsy4-Pro system at a foraging site along the Isar river, Freising, Germany (different day and site than the rest of the data). (**b**) The Clustering analysis of individual calls’ power spectra using PCA and *k*-means clustering, with the optimal number of clusters (*k* = 6) selected via the Davies-Bouldin criterion. The right panel shows the mean power spectrum of each cluster, revealing spectral profiles. (**c**) Example spectrograms of individual calls sampled from each cluster, demonstrating the diversity in spectral shape and frequency content.

#### 2.9.1 Call spectral clustering

To demonstrate the suitability of Batsy4-Pro field recordings for downstream acoustic analyses, an unsupervised spectral clustering approach was applied to extracted echolocation calls. Such analyses are particularly relevant for field datasets, where recordings commonly contain calls from multiple species and a diversity of call types within species. Clustering based on spectral structure provides an objective means of segregating calls before species-focused or behavioural analyses.

For each call, the waveform from the first microphone channel was tapered using a Hann window, and its power spectral density was estimated using MATLAB’s `pspectrum` function with a frequency resolution of 1 kHz. Power spectra were converted to decibel units and truncated to a fixed length of 1024 frequency bins to ensure consistent dimensionality across calls. The resulting spectra were assembled into a matrix representing spectral features for all calls.

Dimensionality was reduced using principal component analysis (PCA), retaining the first ten principal components for clustering. The optimal number of spectral clusters (tested for *k* = 2–6) was determined automatically using the Davies–Bouldin criterion (Davies and Bouldin, 1979) via the `evalclusters` function (MATLAB), providing an objective balance between within-cluster compactness and between-cluster separation. Final clustering was performed using *k*-means on the reduced feature space.

Cluster validity was assessed qualitatively by visualising the mean power spectra for each cluster and by inspecting representative spectrograms randomly selected from each group. This approach illustrates how field-collected data from a compact recorder can be structured into acoustically coherent call groups, facilitating subsequent species identification, call-type classification, or behavioural state analyses without requiring prior labels.

#### 2.9.2 3D localisation via time-difference of arrival

To estimate the spatial position of the calling bat for each detected call, a time-difference-of-arrival (TDoA) approach was applied to the four-channel recordings. Inter-microphone arrival delays were extracted using a custom cross-correlation-based function (`estimateTDOA`) that computed pairwise time lags for all microphone pairs. The resulting delay estimates were passed to a multilateration routine (`localiseTDOA`) that solved for the threedimensional source position (Figure 7) based on the known microphone geometry and an assumed speed of sound of *c* = 343 m s^−1^. These routines form part of the `Array WAH` `– Microphone Array Design Tool` (Umadi, 2025a,d).

**Figure 7.**
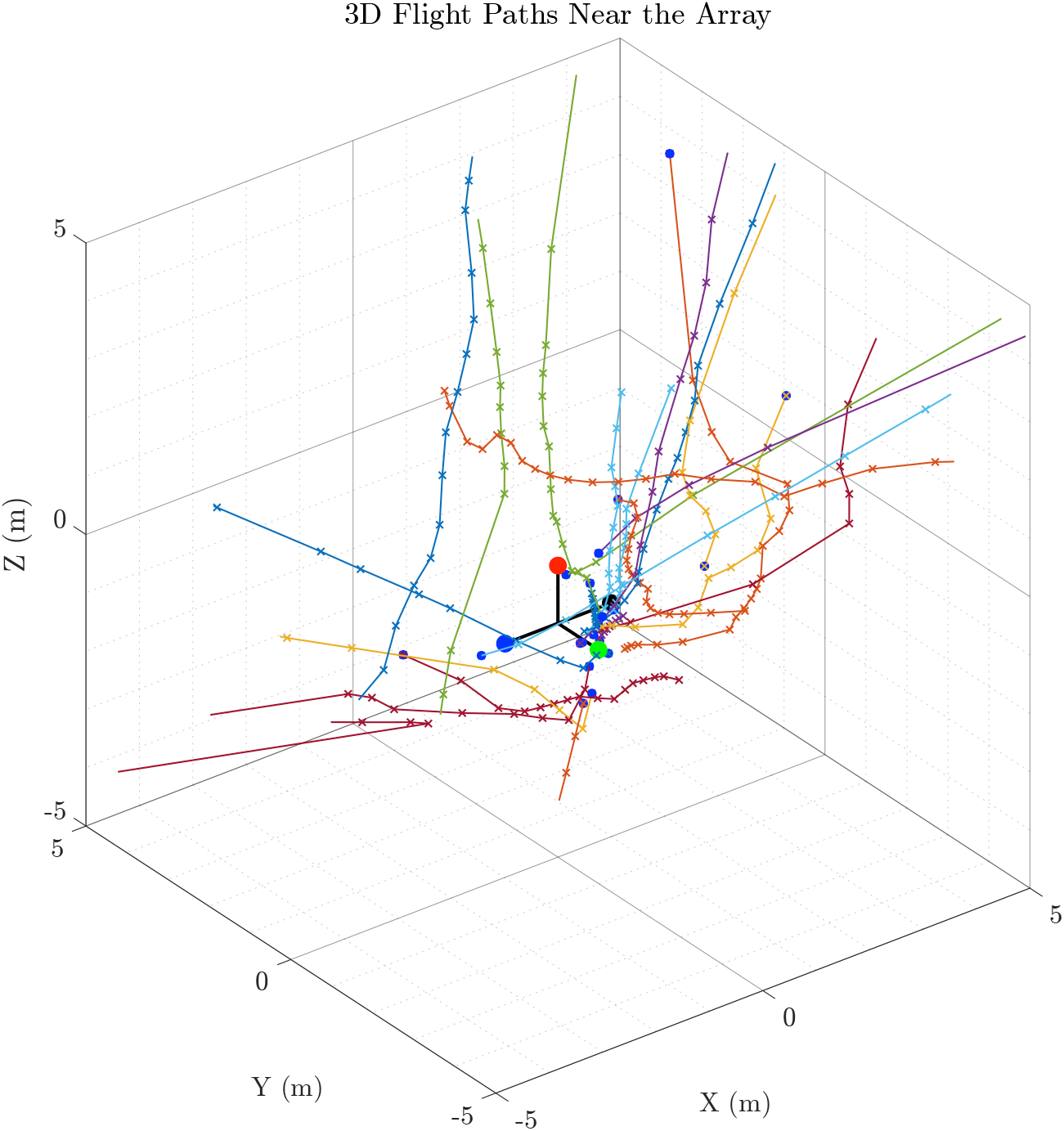
The array was deployed at a bat foraging site, and calls were recorded manually by monitoring the real-time heterodyne output of the system. Using time-difference-of-arrival (TDoA) localisation, the 3D positions of calls were estimated. The flight paths are visualised within ±5 m in all directions for clarity. Microphone positions are shown as large coloured dots (RGBK). The blue dots mark the final call detected in each sequence, corresponding to the last reliably localisable event. The trajectories suggest that bats were actively approaching the array.

### 2.10 Localisation Performance Characterisation

This analysis addressed two deployment-specific questions: where the demonstrated array provides useful localisation performance and how sensitive that performance is to the velocity of a moving source. To answer them, I used the Array WAH forward-evaluation method in a Monte Carlo simulation of the deployed array (Umadi, 2025a,d). The analysis quantified positional uncertainty across the array’s operational volume using a grid-based sampling strategy and included an approximate, velocity-dependent representation of source motion (Figure 8). Here, Array WAH was used to characterise the fixed, deployed geometry; microphone positions were not optimised as part of the present simulation.

**Figure 8.**
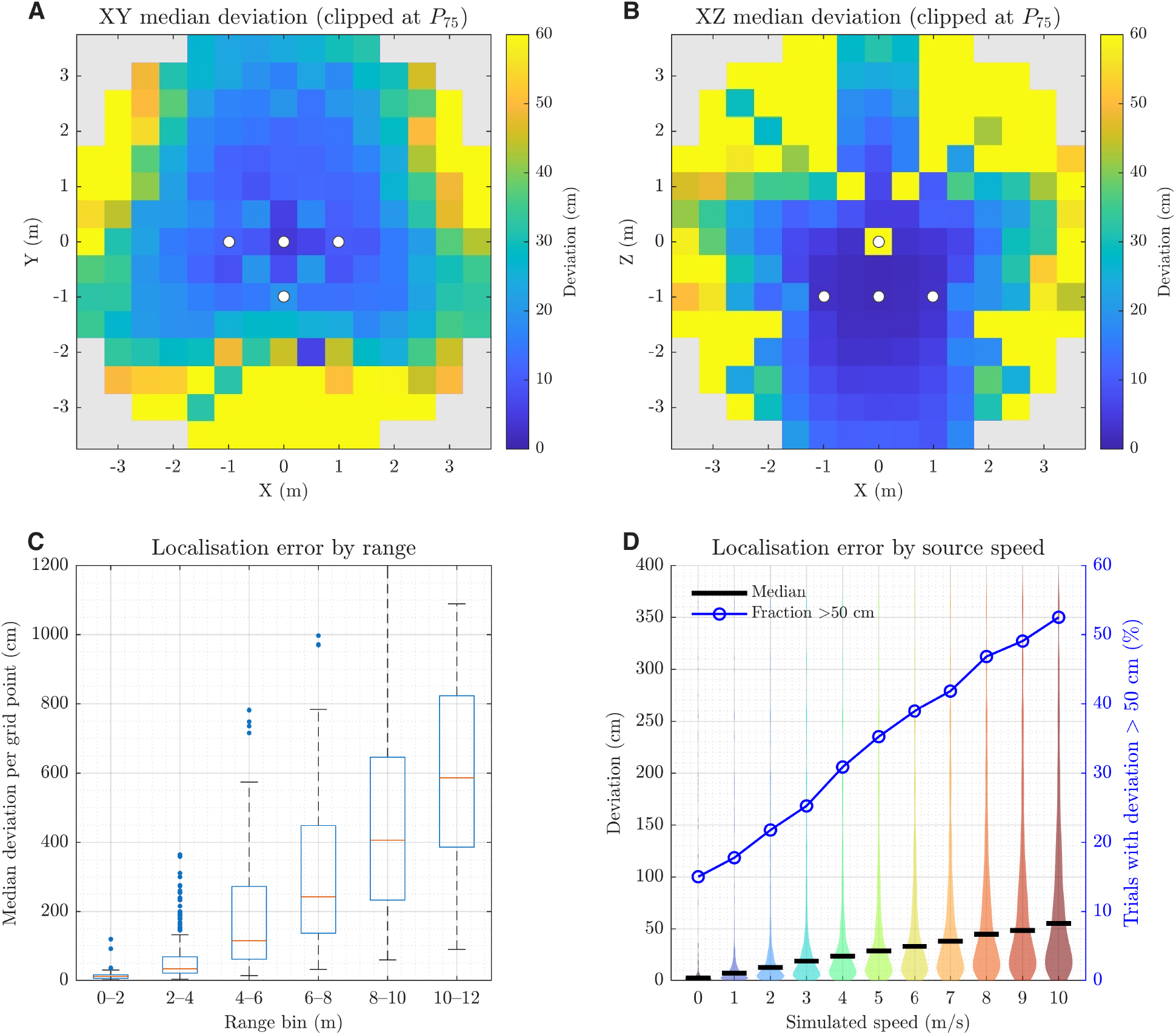
Grid-based localisation error analysis. (**A, B)** Spatial distribution of per-grid-point median localisation deviation in the horizontal XY plane (*z* = 0.5 m) and vertical XZ plane (*y* = 0 m), respectively, for positions less than 4 m from the reference microphone. Values above the 75th percentile (60 cm) are clipped only for colour-scale visualisation and appear at the yellow upper limit. Light-grey cells lie outside the evaluated < 4 m subset and do not represent zero error. White circles indicate microphone positions. **(C)** Per-gridpoint median deviation as a function of reference-microphone range. Boxes aggregate values across simulated speeds and directional trials in 2 m bins over the complete 0–12 m field. **(D)** Trial-wise deviation distributions for simulated speeds of 0–10 m s^−1^ within the <4 m near-field subset. Violin densities are restricted to the displayed 0–400 cm range for legibility, whereas black bars show medians calculated from the complete distributions. The blue line and right axis show the percentage of trials with deviations greater than the fixed 50 cm criterion. Grid resolution: 0.5 m; synthetic call: 5 ms downward quadratic FM sweep (80–25 kHz).

All simulations employed synthetic, downward-sweeping quadratic frequency-modulated calls with a fixed duration of 5 ms (see Section 3.1), spanning 80–25 kHz and generated at 192 kHz. A systematic grid extended ±10 m with 0.5 m spacing. Two intersecting analysis planes were evaluated: a horizontal XY plane at *z* = 0.5 m and a vertical XZ plane at *y* = 0 m. Together, these planes comprised 3362 spatial positions and provided coverage of the near-to mid-field region relevant for field deployments.

Simulated speeds ranged from 0 to 10 m s^−1^ in increments of1m s^−1^. For the stationary condition (*v* = 0 m s^−1^), one trial was simulated per grid point. For every non-zero speed, eight Monte Carlo trials were performed per grid point, with direction randomised in three dimensions. This produced 272,322 attempted trials across all spatial and kinematic conditions.

For each trial, the assigned velocity affected the synthetic call through Doppler resampling and a first-order correction to the microphone-specific propagation delays. The source coordinate used as ground truth remained the grid-point centre; the simulation therefore approximated motion effects rather than explicitly propagating a source along a time-resolved trajectory during call emission. The four microphone signals were rendered into a shared output buffer sized to accommodate the largest relative arrival delay and the complete synthetic waveform in every channel. TDoAs were estimated by cross-correlation, and the source position was recovered by multilateration. Localisation error was quantified as the Euclidean distance between the estimated position and the true grid-point centre, expressed in centimetres (Umadi, 2025d).

Localisation performance was summarised at two levels. Per-grid-point statistics, including median and 95th-percentile deviation, were computed across all speeds and directional trials. At the per-trial level, deviation distributions were analysed as functions of speed and range. Separate univariate linear models were used to calculate the variance in trial-wise localisation error explained by speed and range (*R*^2^). Because the error distributions were strongly non-normal, Spearman rank correlation and a Kruskal–Wallis test were also used to assess the speed association. Range was calculated from the first microphone, which served as the TDoA reference, rather than from the geometric centre of the array. Near-field analyses included positions with this reference-microphone range below 4 m. Of the 272,322 attempted trials, 271,311 returned finite localisation estimates; the 1011 failed trials (0.37%) were excluded from numerical summaries.

## 3 RESULTS

I deployed the Batsy4-Pro system and accompanying fourchannel microphone array during the first week of August 2025 along the river Isar in Freising, Germany (Figure 1). I selected the recording site at the edge of a riparian forest along the riverbank for its consistent bat foraging activity and suitability for demonstrating multichannel ultrasonic recording under field conditions. I mounted the array on a standard photographic tripod and oriented it to face the primary approach corridor used by bats when moving between foraging areas.

Deployment involved three straightforward steps: I as sembled the array arms onto the tripod-mounted hub, connected the microphone cables to the Batsy4-Pro recorder via the single 6-pin connector, and activated the system by connecting a USB power bank. The entire setup process typically requires less than five minutes. The system becomes immediately operational, with the integrated heterodyne monitoring providing real-time auditory feedback of bat activity through headphones.

### 3.1 Data Quality Assessment and Call Selection

I screened recorded call sequences using the *Bat Reviewer* software tool (Umadi, 2025b). The screening process evaluated each call sequence for signal quality, channel consistency, and freedom from contaminating echoes or environmental interference. Of the 30 recorded sequences, I selected 21 for detailed analysis based on the following quality criteria: clear separation between direct calls and ground echoes, consistent detection across all four microphone channels. I retained 368 of the 440 calls (83.6%) for quantitative analysis. This high retention rate indicates robust multichannel recording performance and minimal data loss from technical artefacts.

Call durations exhibited the short, frequency-modulated characteristics typical of aerial-hawking insectivorous bats foraging in semi-open environments. The median call duration was 4.51 ms (interquartile range: 3.64–5.53 ms, *μ* = 4.91 ms), with the central 90% of calls spanning 3.14–9.16 ms. The full range extended from 2.78 ms to 13.26 ms, reflecting variation in call structure associated with different phases of foraging behaviour and target approach distances. Detailed statistics on call duration distributions are provided in Supplementary Figure S1.

I stored all selected calls as individual 20 ms audio clips extracted from the continuous four-channel recordings, preserving synchronised waveforms across all microphone channels. These clips formed the basis for subsequent signal-to-noise characterisation, spectral clustering, and three-dimensional source localisation analyses presented in the following sections.

### 3.2 SNR Characterisation

Across *N* = 368 echolocation calls recorded with Batsy4-Pro, the maximum-channel SNR_peak_ had a median value of 27.3 dB (interquartile range: 10.4 dB; mean ± SD: 27.6 ± 6.6 dB; Fig. 5a). Per-channel SNR_peak_ distributions were broadly similar, with median values ranging from approximately 21 to 26 dB, indicating comparable noise performance across channels.

For comparison, the same analysis was applied to a randomly selected subset of *N* = 500 echolocation calls recorded during a separate experiment dataset using a professional-grade audio interface (Behringer U-PHORIA UMC404HD). The call extraction protocol was the same, with each call corresponding to a 20 ms duration. These data were collected in a different experiment (Umadi and Firzlaff, 2025) using a similarly sized microphone array constructed with the same MEMS microphones, and recorded via a custom MATLAB acquisition script on a MacBook Air (2015). The resulting maximum-channel SNR_peak_ distribution closely matched that obtained with Batsy4-Pro, with a median of 27.4 dB (interquartile range:12.5 dB; mean ± SD: 29.0 ± 9.9 dB; Fig. 5b). Together, these results indicate that variability in SNR is dominated by differences in call amplitude rather than fluctuations in recorder noise, and that the field performance of Batsy4-Pro approaches that of a professional soundcard-based recording system.

### 3.3 Characterising Expected Localisation Performance

Within 4 m of the reference microphone, 386 grid points and 31,151 valid trials were evaluated. The spatial distribution of median localisation error exhibited pronounced structure across both horizontal and vertical planes (Figure 8). The lowest errors were concentrated around the array, while larger errors occurred preferentially towards the edge of the evaluated region and in geometrically unfavourable directions.

This spatial pattern was consistent across planes and was not well described by distance alone. Instead, uncertainty varied substantially among grid points at comparable ranges, reflecting anisotropic conditioning imposed by the array geometry. Positions that subtended favourable angular separations relative to multiple microphone baselines exhibited high localisation precision, whereas positions aligned with extended baselines or producing weak geometric dilution showed degraded performance, consistent with the results from Umadi (2025d).

The distribution of per-point median deviations increased with range. In the 0–2 m bin, the median was 11.5 cm (interquartile range: 5.7–16.0 cm); in the 2–4 m bin, it was 33.4 cm (20.8–67.8 cm). Across all 386 near-field grid points, the median per-point error was 27.3 cm (14.1–59.5 cm). The full-field range panel further shows the rapid deterioration outside this near-field region.

Within the near-field region, 71.0% of grid points (274 of 386) achieved median localisation deviations of 50 cm or less, and 84.7% (327 of 386) achieved deviations of 100 cm or less.

Performance declined between the two near-field range bins. In the 0–2 m bin, 94.4% of grid points (85 of 90) achieved median deviations ≤ 50 cm and 96.7% (87 of 90) achieved ≤ 100 cm. In the 2–4 m bin, the corresponding proportions were 63.9% (189 of 296) and 81.1% (240 of 296), respectively. The substantial variation among positions within each bin demonstrates that range alone does not capture the spatially anisotropic error pattern.

#### Sensitivity of simulated localisation error to source speed

**T**he implemented motion approximation produced a systematic shift towards larger localisation errors as simulated speed increased. Median trial-wise error rose from 2.18 cm in the stationary condition to 55.28 cm at 10 m s^−1^. Over the same interval, the interquartile range changed from 0.75–12.24 cm to 20.63–128.76 cm.

The fraction of trials exceeding 50 cm increased monotonically from 15.0% at 0 m s^−1^ to 52.5% at 10 m s^−1^. Speed was positively associated with error by both Pearson correlation (*r* = 0.028, *p* < 0.001) and Spearman rank correlation (*ρ* = 0.341, *p* < 0.001), and error distributions differed among speed conditions (Kruskal–Wallis test, *p* < 0.001). A linear model fitted to the raw, strongly right-skewed trialwise errors nevertheless yielded a small *R*^2^ of 0.0008 for speed; the corresponding range-only model yielded *R*^2^ = 0.0116.

The difference between the low linear-model *R*^2^ and the clearer rank-based and distributional trends reflects the highly skewed, heteroscedastic error distributions and the presence of large geometric outliers. Accordingly, the simulated speed effect is better described by changes in medians, quantiles, and threshold exceedance than by the fraction of raw-error variance explained by a linear model.

These velocity-dependent results must be interpreted within the scope of the simulation. Motion was represented through Doppler resampling and a first-order delay correction at a fixed source coordinate, rather than through explicit time-resolved propagation of a moving source. The analysis therefore establishes sensitivity of this implementation to the velocity parameter but does not by itself quantify the accuracy expected for real moving animals.

## 4 DISCUSSION

The development of Batsy4-Pro addresses a persistent constraint in behavioural bioacoustics: the trade-off between recording quality, portability, and cost. Traditional multichannel setups for studying echolocating bats typically require a combination of professional audio interfaces, external preamplifiers, laptop computers, and reliable power infrastructure, and have been used in lab and field-based studies (Brinkløv et al., 2011; de Framond et al., 2023; Eitan et al., 2022; Gaudette and Simmons, 2014; Jakobsen et al., 2025; Kounitsky et al., 2015; Surlykke et al., 2009; Umadi and Firzlaff, 2025; Xia et al., 2025) (also see (Koblitz, 2018)). These systems, while capable of excellent signal fidelity, present significant operational challenges in field settings. Proper channel configuration, gain staging, weatherproofing sensitive components against condensation, and power management require both technical expertise and logistical support, which necessitate team-based deployments. The cumulative cost of such configurations typically ranges from several hundred to several thousand euros, compounding these deployment constraints.

Batsy4-Pro offers a fundamentally convenient approach by integrating all essential components – analogue-to-digital conversion, computational processing, data storage, and real-time monitoring – into a single compact device weighing less than 150 grams. The system requires no external laptop, no multichannel soundcard, and no AC power source, operating instead from a standard USB power bank. This architectural consolidation eliminates the multiplicity of failure points inherent in distributed recording chains, dramatically simplifying field deployment. A researcher can carry the complete system in a backpack, assemble the array in minutes, and begin recording without the need for calibration procedures or driver configuration that plague computer-based acquisition systems.

The baseline calibration of the system represents another significant advantage. Because all four channels share identical signal paths—from MEMS microphone through ADC to digital storage—inter-channel gain differences are minimised by design rather than requiring *post-hoc* correction. Unlike audio interfaces where each channel may have independent gain controls that drift or reset between sessions, Batsy4-Pro maintains consistent relative sensitivity across channels. Level calibration to an absolute acoustic reference pressure (20 *μ*Pa) needs to be performed only once, and the resulting compensation factors can be incorporated directly into the firmware or applied during post-processing. This inherent calibration stability is particularly valuable for studies requiring long-term projects or comparative measurements across multiple recording sessions.

For applications demanding higher precision, frequency-response calibration filters can be computed using established methods (Barrera-Figueroa, 2018; MacDonald and Tran, 2007; *IEC 61094-1* 2000; *IEC 61094-3* 2016; Wathen-Dunn, 1949) and integrated into the processing pipeline. Given the computational capability of the Teensy 4.1 platform, such filtering can be applied in real time before data are written to storage, analogous to the heterodyne downconversion already implemented in the current firmware. This flexibility to incorporate domain-specific signal conditioning at the firmware level—rather than relying on postprocessing workflows—represents a significant advantage for researchers working with non-standard microphone arrays or specialised acoustic environments.

### 4.1 Cost Accessibility and Open Development Model

The total component cost for Batsy4-Pro, including the recorder unit and four-channel array, is approximately 200 euros when sourced in small quantities from local suppliers. This represents less than one-tenth the cost of comparable commercial multichannel ultrasonic recording systems, or using a professional audio interface with a computer (Table 1).

Importantly, all hardware designs, firmware source code, and assembly documentation are released as open source under the CERN Open Hardware Licence v2 – Strongly Reciprocal (CERN-OHL-S), GPLv3, and CC-BY-SA-4.0 licences, respectively (Umadi, 2026b). These licences explicitly permit commercial manufacture, distribution, and sale without royalty obligations, provided that derivative works remain openly licensed and complete source documentation is made available to recipients. This deliberate choice encourages local manufacturing initiatives, institutional workshop projects, and community-driven development, thereby further reducing costs and increasing accessibility.

Lower component cost and open licensing address only the equipment dimension of accessibility. Effective adoption also depends on component availability, fabrication facilities, documentation, technical support, and training.

Research on echolocation in bats and other animals is inherently interdisciplinary, drawing on concepts and tools from electronics, signal processing, and communication engineering, as well as biological theory. Since the earliest demonstrations of ultrasonic biosonar in bats, progress in this field has depended on close collaboration between biologists and engineers. Notably, Donald Griffin and Robert Galambos’s foundational work on bat echolocation was enabled by access to specialised ultrasonic detection equipment developed by the physicist G. W. Pierce at Harvard (Griffin, 1958, 2001; Griffin and Galambos, 1941; Pierce and Griffin, 1938). This pattern has persisted: advances in understanding biosonar behaviour, sensory-motor coordination, and ecological function have consistently required custom instrumentation and bespoke analytical methods.

While scientifically productive, this reliance on engineering expertise can pose substantial barriers for biologists and ecologists, particularly when specialised hardware is expensive, difficult to deploy, or available only in a small number of laboratories. Recent developments in lightweight acoustic tags and on-animal recorders have demonstrated how accessible instrumentation can transform the study of natural behaviour, enabling unprecedented insights into free-flight echolocation, foraging strategies, and sensorimotor control (Stidsholt et al., 2025a; Stidsholt, 2025; Stidsholt et al., 2019, 2021, 2025b). Similarly, neuroethological studies employing custom-developed wireless neural telemetry systems have revolutionised our understanding of spatial navigation by enabling single-neuron recordings from bats flying in naturalistic environments—from large indoor flight rooms to 200-meter tunnels and even on remote oceanic islands (Eliav et al., 2021; Palgi et al., 2025; Yartsev and Ulanovsky, 2013). However, such technologies often remain closely coupled to the groups that develop them, limiting broader adoption and replication.

By providing a complete, tested, and documented design that can be assembled from globally available components, Batsy4-Pro lowers the threshold for entry into experimental bioacoustics. The choice to base the system on the widely adopted Teensy platform, supported by extensive community documentation and a large user base among hobbyists and engineers, ensures that technical support and troubleshooting resources are readily available outside traditional academic networks. This accessibility extends beyond professional researchers to educators, citizen scientists, and conservation practitioners who may wish to incorporate behavioural monitoring into field programmes but lack access to expensive commercial systems.

### 4.2 Integrated Real-Time Monitoring and Adaptive Recording

The methodological value of integrated real-time monitoring lies in coupling immediate auditory assessment with event-triggered recording.

Researchers can listen for specific acoustic signatures—such as approach sequences, social calls, or feeding buzzes—and selectively capture relevant events rather than record continuously and screen long recordings post hoc. The retrospective buffer complements this process by preserving the five seconds preceding a trigger, retaining the behavioural context before the decision to record.

The tunable 10–85 kHz carrier permits rapid monitoring across call bands when multiple species or call types are present. Direct passthrough extends monitoring to audible-frequency applications, while the combined-input mode permits simultaneous auditory oversight of all four microphone positions.

### 4.3 Signal Quality and Localisation Performance

The signal-to-noise characterisation presented in this study demonstrates that Batsy4-Pro achieves recording fidelity comparable to professional-grade audio interfaces under field conditions. Median maximum-channel SNR values of 27.3 dB across 368 echolocation calls, with consistent performance across all four channels, indicate that the system noise floor is sufficiently low to support high-quality acoustic analysis. The close correspondence between SNR distributions obtained with Batsy4-Pro and those from a commercial soundcard-based system (median 27.4 dB, *N* = `500` calls) confirms that observed SNR variability is dominated by natural variation in call amplitude and source distance rather than by recorder noise characteristics. This performance supports the use of compact MEMS microphones and integrated ADC architectures for demanding bioacoustic applications that previously required more complex recording chains.

The spatial localisation simulations provide a quantitative basis for interpreting positional estimates derived from the four-channel cross-type array configuration employed here. Within 4 m of the reference microphone, 71.0% of grid points achieved median localisation errors no greater than 50 cm and 84.7% no greater than 100 cm. Performance was strongest within 2 m, where 94.4% of positions met the 50 cm criterion, and declined with range in a spatially anisotropic pattern. These results delineate a useful near-field volume for behavioural studies focused on close-range interactions, such as prey capture, obstacle avoidance, or social encounters.

The simulations also showed a velocity-dependent shift in the error distributions: median trial-wise error increased from 2.18 cm in the stationary condition to 55.28 cm at 10 m s^−1^, and the fraction of errors greater than 50 cm increased from 15.0% to 52.5%. This pattern indicates that the localisation pipeline is sensitive to the velocity-dependent Doppler resampling and delay correction implemented in the simulation. It does not, however, demonstrate the magnitude of motion-related error expected in the field because source movement was approximated at a fixed grid coordinate rather than propagated explicitly throughout the call.

The very small linear-model *R*^2^ for speed should therefore not be interpreted as evidence that the distributions were invariant. The monotonic changes in median error, rank correlation, and 50 cm threshold exceedance all show a distributional shift, whereas the raw-error regression was dominated by strong skew, heteroscedasticity, and large geometry-dependent errors. Reporting several complementary summaries is consequently more informative than relying on a single measure of variance explained.

These findings motivate further validation with a physically explicit moving-source model and controlled playback trajectories. Such work should update microphone-specific source distance continuously during emission and use radial source–receiver velocity for Doppler calculations. Until that validation is available, the stationary and spatial-grid results provide the firmer basis for defining the array’s usable volume, while the velocity analysis should be treated as a model-sensitivity assessment.

The present array geometry was selected for portability and rapid deployment rather than numerically optimised for extended tracking range. The increasing uncertainty beyond 4 m illustrates the geometric trade-off between portability and spatial coverage. WAH-*i* provides a route for investigating task-specific optimisation of the four-microphone geometry, as considered below (Umadi, 2026f).

### 4.4 Validation Against Established Methods and Ground Truth Considerations

The localisation performance presented here must be interpreted in the context of broader methodological challenges in bat bioacoustics. Many published studies employing microphone arrays with 8–12 channels report localisation results primarily for the final approach to terminal phase (<2 s) of echolocation sequences, where signal-to-noise ratios are highest and geometric configurations most favourable (Jakobsen et al., 2025; Xia et al., 2025). Screening procedures that exclude spatial jumps, far-field data points, and low-SNR calls are routinely applied, yet the underlying ground-truth accuracy of these systems is rarely quantified or reported in detail. Instead, reliability is inferred indirectly from signal quality metrics and trajectory smoothness, with the implicit assumption that high-SNR calls localise accurately. Channel count alone is therefore not a sufficient proxy for spatial performance. Under the fixed-aperture conditions examined with WAH-*i*, optimised four-microphone arrays matched or exceeded the volumetric localisation performance of higher-order arrays with suboptimal geometries, although these outcomes remain conditional on the chosen observation volume, signal model, accuracy criterion, and physical constraints (Umadi, 2025d, 2026f).

This lack of standardised validation against known source positions represents a significant gap in the field. While simulation-based approaches provide valuable estimates of expected performance under idealised conditions, empirical validation using calibrated acoustic sources at surveyed positions remains essential for establishing absolute accuracy. The discrepancy between simulated and empirical accuracy is likely to be non-negligible, particularly in field environments where multipath propagation, atmospheric absorption, and environmental noise degrade signal quality in ways that are difficult to model comprehensively. However, the results are consistent with empirical validation studies such as de Framond et al. (de Framond et al., 2023) – a rare example of explicit ground-truth validation – who assessed localisation accuracy using calibrated loudspeaker playbacks at 45 precisely surveyed positions around a fixed array and reported strong spatial dependence of error.

The modular Batsy array facilitates this validation by allowing candidate geometries to be assembled, surveyed, and tested using calibrated playback from known positions. Predicted performance can therefore be compared directly with the accuracy of the realised configuration under field-relevant acoustic conditions.

### 4.5 Geometry-optimised extensibility and future development directions

Extensibility in a multichannel recorder is commonly interpreted as an increase in channel count. For acoustic localisation, however, adding microphones does not necessarily produce a proportional improvement if the resulting geometry is poorly conditioned for the region of interest. WAH-*i* offers a complementary form of extensibility by optimising the placement of the four microphones already supported by Batsy4-Pro . Rather than assuming a universally optimal array, the optimisation is defined by the biological task: the intended three-dimensional observation volume, an acceptable localisation-error threshold, the acoustic signal and noise model, and physical constraints such as maximum aperture and minimum inter-microphone separation (Umadi, 2026f). The resulting configuration is consequently a task-specific Batsy array rather than a replacement recorder architecture.

WAH-*i* exports the optimised microphone geometry as Cartesian coordinates, with one row per microphone and positions expressed in metres relative to the array centroid (Umadi, 2026e). For a mounting system referenced to a common origin, the coordinate of microphone *i*, p_*i*_ = (*x*_*i*_, *y*_*i*_, *z*_*i*_), can be translated into radial distance, azimuth, and elevation as

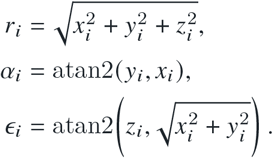

A reproducible conversion requires an explicit coordinate convention; for example, +*x* may point towards the centre of the observation volume, +*z* vertically upwards, and positive azimuth from +*x* towards +*y*. If individual arm pivots do not share the coordinate origin, the corresponding pivot offset must be subtracted before calculating these settings. The detailed engineering of a reconfigurable mounting hub lies beyond the scope of the present study, but its essential design considerations follow directly from the conversion: sufficient radial reach, azimuth and elevation range, known graduation increments, mechanical rigidity, minimum microphone clearance, and independent control of microphone orientation.

The conversion from a tetrahedral starting configuration to an example WAH-*i* geometry, including the radial and angular adjustment required for each microphone, is illustrated in Supplementary Figure S2.

The finite resolution of a physical mounting system can be incorporated into the same workflow. The WAH-*i* values of *r*_*i*_, *α*_*i*_, and *ϵ*_*i*_ can first be rounded to the available radial and angular increments. The mechanically realisable coordinate can then be reconstructed as

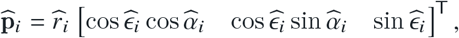

and the reconstructed geometry returned to the simulation framework. This closes the computational–physical loop by quantifying any loss of predicted performance caused by angular graduation, arm-length increments, or other mechanical constraints before field deployment. After assembly, the microphone positions should be surveyed and the measured coordinates used for a final simulation and for subsequent TDoA localisation. Controlled playback from known positions would then test whether the realised array achieves the predicted operating volume.

WAH-*i* thus provides task-specific geometry design for the four-channel system, while Batsy supplies the portable acquisition platform needed to realise and test that geometry. Increasing the electronic channel count remains valuable where four microphones cannot meet the required coverage or robustness, but it becomes a requirement demonstrated by task-level analysis rather than a default design assumption.

The architecture of Batsy4-Pro is inherently extensible in both hardware and software dimensions. The Teensy 4.1 platform supports dual I^2^S interfaces capable of handling up to eight synchronous audio channels with appropriate ADC configurations, and time-division multiplexing (TDM) protocols can extend this to sixteen channels for applications requiring higher spatial resolution or increased angular coverage. While TDM implementation is beyond the scope of the current work, the underlying microcontroller capabilities and firmware structure are compatible with such expansion, requiring modifications primarily to the I^2^S configuration and buffer management routines.

Sampling rate extensibility represents another avenue for development. Although the current implementation operates at 192 kHz—adequate for most bat species and ultrasonic bioacoustic applications—certain taxa produce calls with substantial energy above 120 kHz, including many rhinolophid and hipposiderid species. Higher sampling rates (384 kHz or 768 kHz) can be achieved using alternative ADC hardware, such as the AK5572 from Kaamos Tech (Nihtila, 2023), which offers dual-channel 32-bit conversion at up to 768 kHz. Similarly, while the current system employs 16-bit resolution to maintain compatibility with the *Teensy Audio Library* and minimise memory requirements, the WM8782 ADCs support 24-bit conversion. Implementing 24-bit recording would require custom firmware that bypasses the Audio Library’s 16-bit constraints, but such increased bit depth is unlikely to yield substantial practical improvements for most field applications, where environmental noise rather than quantisation noise determines the effective signal-to-noise ratio.

Beyond audio acquisition, the Teensy 4.1 provides extensive support for auxiliary sensors via I^2^C, and serial interfaces. Integration of GPS modules for geolocation and precision timestamping, environmental sensors (temperature, humidity, barometric pressure) for atmospheric correction, passive infrared or photogate sensors for motion-based triggering, and anemometers for wind monitoring all represent straightforward extensions that can enhance the contextual richness of acoustic datasets. Such multimodal data streams are particularly valuable for behavioural ecology, where understanding the environmental conditions and physical constraints under which animals operate is essential for interpreting acoustic behaviour.

### 4.6 Broader Implications for Behavioural Research and Conservation

The availability of low-cost, field-ready multichannel ultrasonic recording systems has implications that extend beyond individual research projects. By reducing the practical burden of multichannel acquisition, such systems can support experiments across a broader range of species, habitats, and behavioural contexts.

Recent calls to incorporate behavioural diversity into biodiversity monitoring highlight the value of tools that extend monitoring beyond species detection (Mikula et al., 2026); by combining synchronised acoustic recording with source localisation, Batsy4-Pro can support both targeted behavioural experiments and conservation-oriented monitoring.

Tropical and subtropical regions, which harbour much of the world’s bat diversity and include critical areas for conservation, remain underrepresented in behavioural bioacoustics (Van Helden, 2012). Instrumentation is only one contributor to this imbalance, alongside funding, institutional capacity, training, and field logistics. By reducing acquisition costs and supporting local assembly, Batsy4-Pro may help address one practical barrier, but it should be understood as an enabling component rather than a solution to the broader geographic disparity.

Recent work using Batsy4-Pro to investigate spectral ripple structure in bat echolocation sequences (Umadi, 2026d) demonstrates how such portable systems enable direct empirical testing of ecological and sensory theories under natural field conditions. The responsivity theory provides a process-based analytical and diagnostic framework for generating testable hypotheses about echolocation timing during prey pursuit and interception; coupled with task-specific geometry optimisation through WAH-*i*, Batsy4-Pro can supply the spatially resolved field recordings needed to test those hypotheses and realise the experimental paradigms proposed by the framework (Umadi, 2026f; Umadi and Firzlaff, 2025).

Educational and citizen science applications represent another domain where accessible ultrasonic recording technology can have a significant impact. Current single-channel bat detectors and recorders have proven popular among amateur naturalists, educators, and conservation volunteers, fostering engagement with chiropteran ecology and building public support for bat conservation initiatives. Enabling behavioural experiments—rather than solely passive monitoring—in educational contexts can deepen this engagement and contribute to scientific literacy. Students and citizen scientists equipped with multichannel recording systems can participate meaningfully in research projects, collecting high-quality data that complements professional investigations and expands the geographic and temporal scope of acoustic monitoring efforts.

## 5 CONCLUSIONS

Batsy4-Pro provides an open-source, portable four-channel platform for synchronised ultrasonic recording, selectable real-time monitoring, and three-dimensional acoustic localisation. Field recordings and controlled analyses demonstrate signal quality suitable for behavioural bioacoustics, while the grid simulations characterise a useful near-array operating volume and expose geometry-dependent limits. Coupling the recorder with task-specific array optimisation through WAH-*i* offers a practical route to improving four-channel localisation without necessarily requiring increased channel count. Together, these capabilities can enable novel field experiments and open new avenues for a broader comparative understanding of echolocation behaviour across bat taxa.

## DECLARATIONS

### Ethics approval and consent to participate

The study used only passive, non-invasive acoustic recordings of free-flying wild bats. No animals were captured, handled, restrained, marked, or subjected to experimental manipulation. Ethical approval and consent to participate were therefore not required under applicable local and institutional guidelines, and no permits were required for passive acoustic recording at the study site.

### Consent for publication

Not applicable.

### Availability of data and materials

The datasets generated and/or analysed during the current study, together with the analysis code and firmware for the described system, are available in the Zenodo repository at https://doi.org/10.5281/zenodo.22791790 (Umadi, 2026a).

### Authors’ contributions

The sole author conceived and designed the study, developed the hardware, firmware, and analysis software, collected and analysed the data, performed the simulations, prepared the figures, and wrote and revised the final manuscript.

#### Acknowledgements

I thank Uwe Firzlaff for valuable discussions and constructive feedback during the development of the recording system, and Christian Fink for his assistance with the machining and assembly of the microphone array. I am sincerely grateful to the anonymous reviewers for their insightful engagement with the study and constructive feedback, which made the final manuscript a clearer and more coherent account of the work.

## REVISION SUMMARY

The manuscript revisions are grouped below by public manuscript version.

### Version 2 – peer-review revision

The manuscript was substantially revised and expanded in response to the editor and reviewers’ comments, with the overarching goal of improving accessibility for a broader bioacoustics readership while strengthening performance validation and contextualisation. Key revisions included:

1. Major reorganisation of the system description, now led by a hardware/firmware architecture and signal-flow overview.
2. Addition of a terminology table defining acronyms and embedded audio concepts in accessible language, and a table with comparison to prominent commercially available/open source recorders for contrasting placement of novelty.
3. Inclusion of two dedicated flowcharts describing the system architecture and full recording sequence, explicitly linking the two firmware-defined capture modes (*TAP*: retrospective buffer dump; *HOLD*: buffered-forward recording with prefill) to the ring-buffer and WAV writing steps.
4. Expansion of Methods/Results to quantitatively support core performance claims, including SNR characterisation and a grid-based Monte-Carlo localisation uncertainty analysis using the Array WAH toolkit and Widefield Acoustics Heuristics.
5. A substantially expanded Discussion and comparative table situating the present system against representative commercial and open-source systems, clarifying novelty, intended use-cases, and extensibility.

Further, the manuscript now clarifies that Batsy4-Pro has obtained OSHWA certification (DE165), providing externally validated openness standards. Hardware design files and firmware are released under permissive licenses (CERN-OHL-S, GPLv3).

### Version 3 – public preprint update

(i) Correction and complete reanalysis of the grid-based localisation simulation using delay-aware signal buffers that retain the complete waveform at every microphone. The corrected analysis revealed a sharp boundary in useful localisation performance at approximately 4 m; the associated text and Figure 8 were updated accordingly.

### Version 4 – revision following peer review

The manuscript and associated open-source materials were refined following detailed feedback from two reviewers. Key revisions included:

i. Clarification of why the array simulations are integral to the recorder’s intended use in behavioural field experiments: they define the spatial operating field of the demonstrated geometry and examine how source movement affects localisation performance.
ii. Expansion of reproducibility information, including the exact sampling rate and theoretical frequency limit, measured microphone positions, component-to-component connections, microphone replacement considerations, and the operation and engineering rationale of the buffered recording modes.
iii. Extension of the real-time monitoring firmware and user interface to provide rotary selection of heterodyne carrier frequency, microphone channel, and output level, together with direct passthrough (PT) and four-channel mixed monitoring (MX). The system and ring-buffer diagrams were updated to reflect the implemented signal paths and controls.
iv. Revision of the abstract, captions, Methods, and Discussion for greater precision and accessibility, with additional recent literature placing the system in the context of behavioural bioacoustics, biodiversity monitoring, and comparable multichannel recording approaches.

## SUPPLEMENTARY INFORMATION

## A COMPONENT CONNECTIONS

Table T1 records the signal-level connections used in the reported build. Module manufacturers sometimes use different labels for the same signal; connections should therefore be made by signal function rather than by the physical ordering of pins on a particular breakout board. All modules share a common ground, and supply voltage must follow the specification of the particular module used.

**Table T1.**
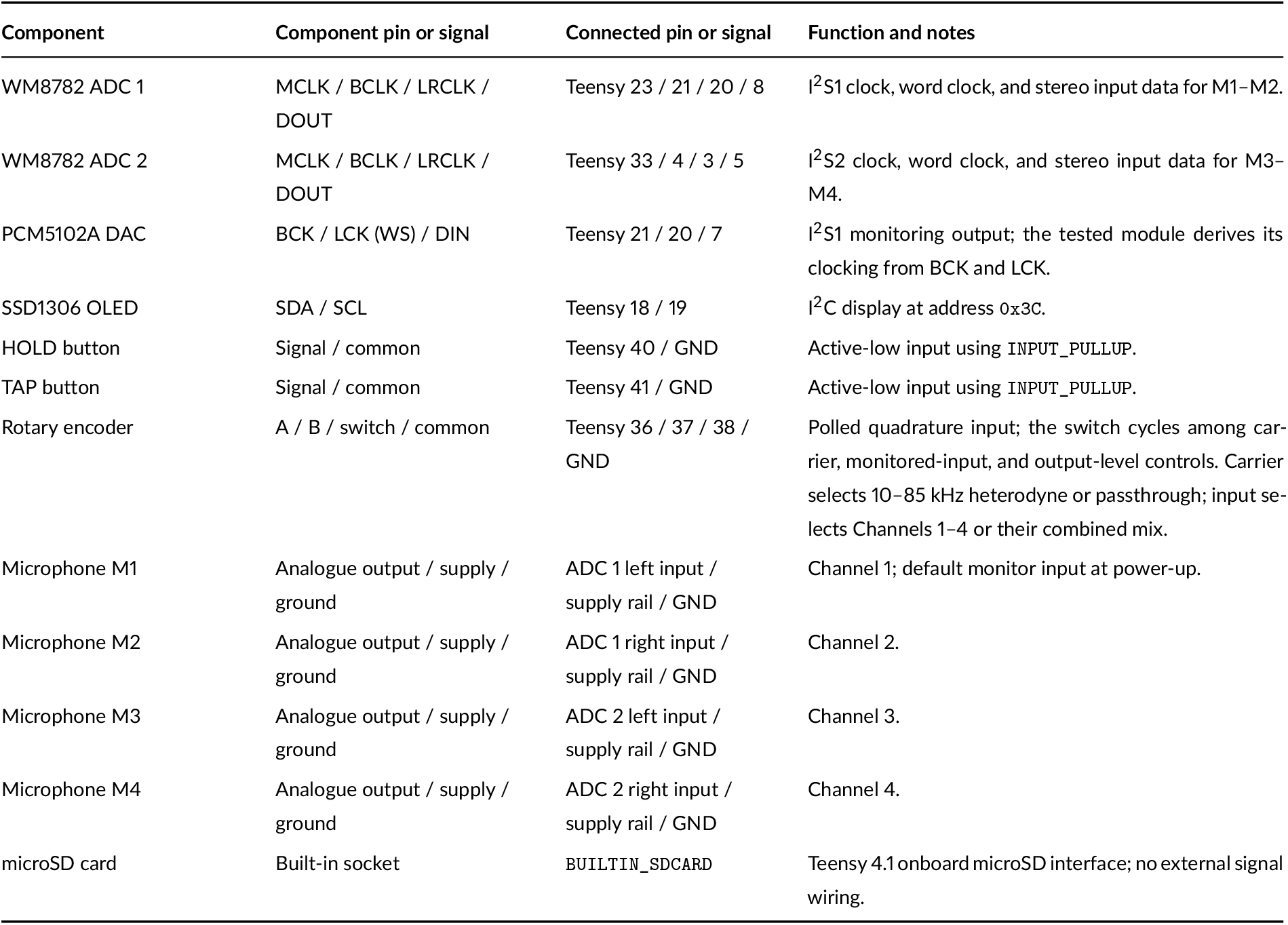
Pin-to-pin connections for each component in the reported Batsy4-Pro build. The two WM8782 boards use separate Teensy I^2^S input interfaces. The PCM5102A shares the I^2^S1 clocks and receives the processed monitoring signal on the Teensy transmit-data line.

| Component | Component pin or signal | Connected pin or signal | Function and notes |
| --- | --- | --- | --- |
| WM8782 ADC 1 | MCLK / BCLK / LRCLK / DOUT | Teensy 23 / 21 / 20 / 8 | I<sup>2</sup>S1 clock, word clock, and stereo input data for M1–M2. |
| WM8782 ADC 2 | MCLK / BCLK / LRCLK / DOUT | Teensy 33 / 4 / 3 / 5 | I<sup>2</sup>S2 clock, word clock, and stereo input data for M3–M4. |
| PCM5102A DAC | BCK / LCK (WS) / DIN | Teensy 21 / 20 / 7 | I<sup>2</sup>S1 monitoring output; the tested module derives its clocking from BCK and LCK. |
| SSD1306 OLED | SDA / SCL | Teensy 18 / 19 | I<sup>2</sup>C display at address 0x3C. |
| HOLD button | Signal / common | Teensy 40 / GND | Active-low input using INPUT_PULLUP. |
| TAP button | Signal / common | Teensy 41 / GND | Active-low input using INPUT_PULLUP. |
| Rotary encoder | A / B / switch / common | Teensy 36 / 37 / 38 / GND | Polled quadrature input; the switch cycles among carrier, monitored-input, and output-level controls. Carrier selects 10–85 kHz heterodyne or passthrough; input selects Channels 1–4 or their combined mix. |
| Microphone M1 | Analogue output / supply / ground | ADC 1 left input / supply rail / GND | Channel 1; default monitor input at power-up. |
| Microphone M2 | Analogue output / supply / ground | ADC 1 right input / supply rail / GND | Channel 2. |
| Microphone M3 | Analogue output / supply / ground | ADC 2 left input / supply rail / GND | Channel 3. |
| Microphone M4 | Analogue output / supply / ground | ADC 2 right input / supply rail / GND | Channel 4. |
| microSD card | Built-in socket | BUILTIN_SD CARD | Teensy 4.1 onboard microSD interface; no external signal wiring. |

## B I^2^S CLOCK SETTING

**Listing 1.**
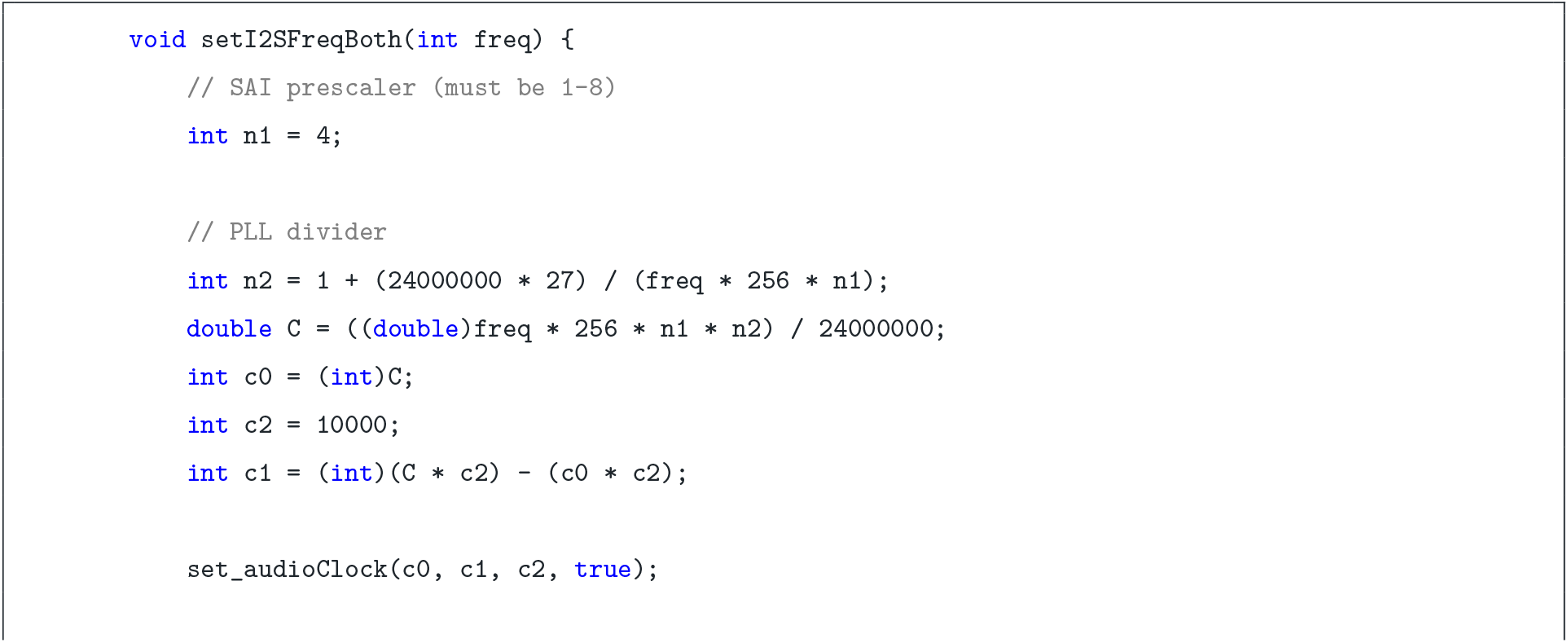
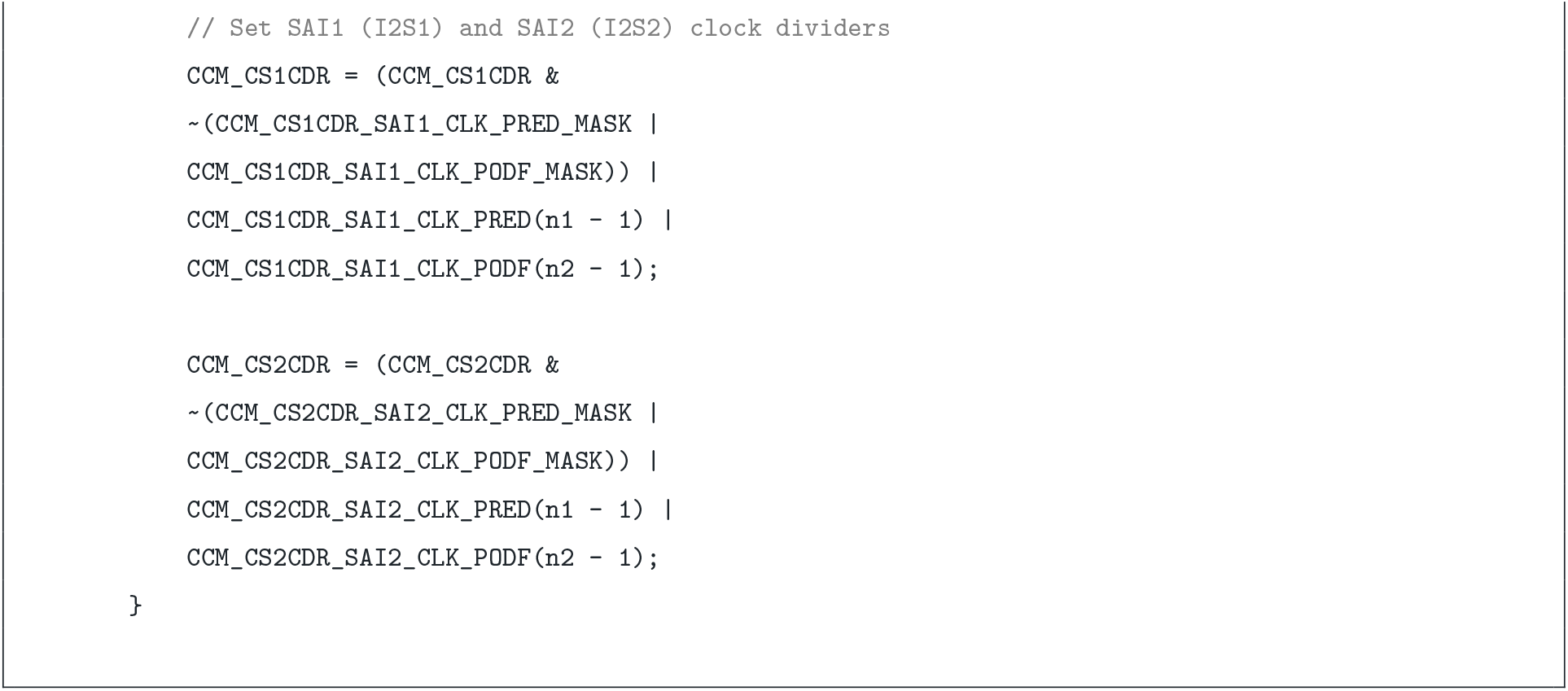
^2^S clock configuration for 192 kHz sampling on Teensy 4.1.

## C RING BUFFER IN PSRAM

**Listing 2.**
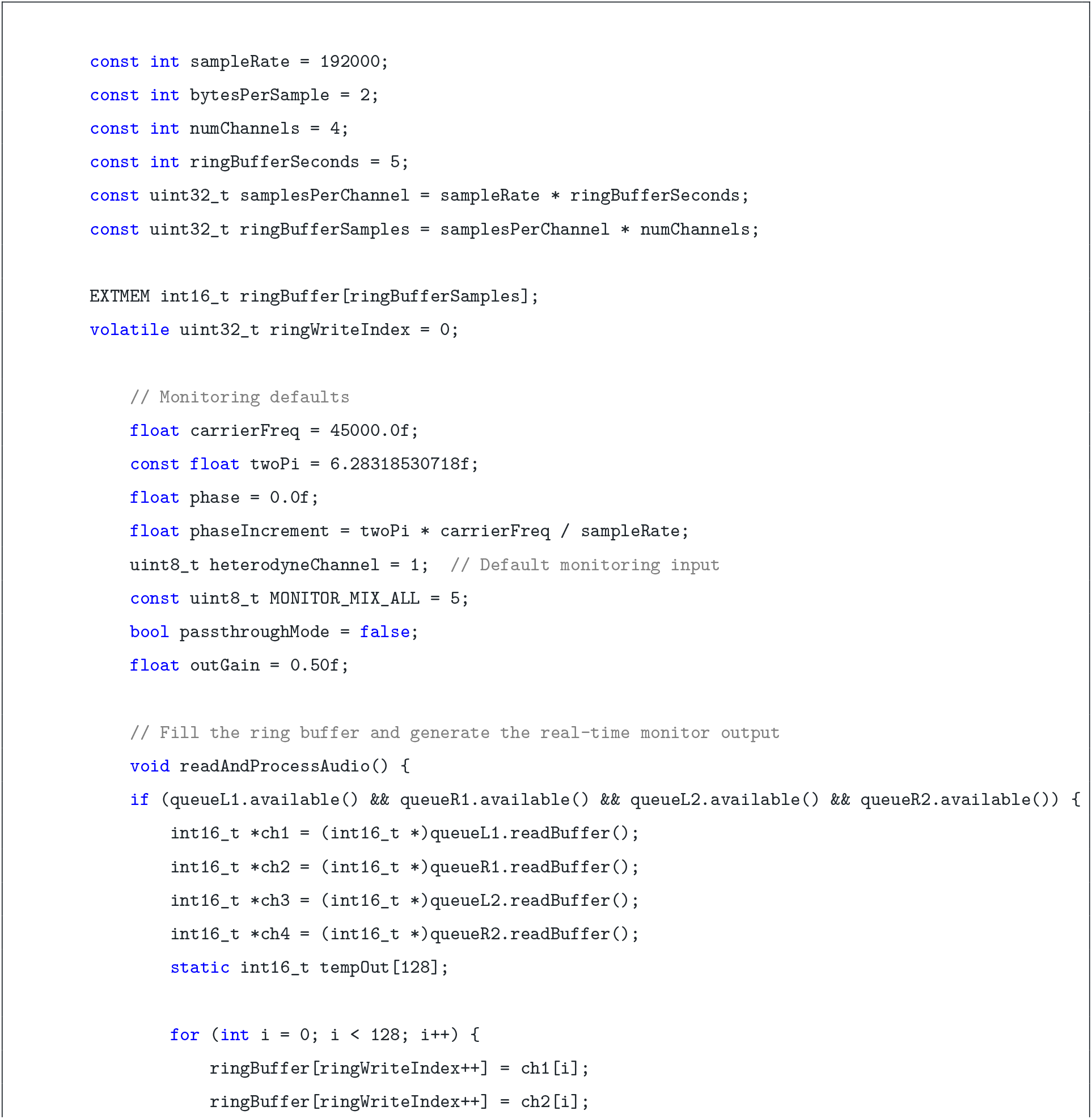
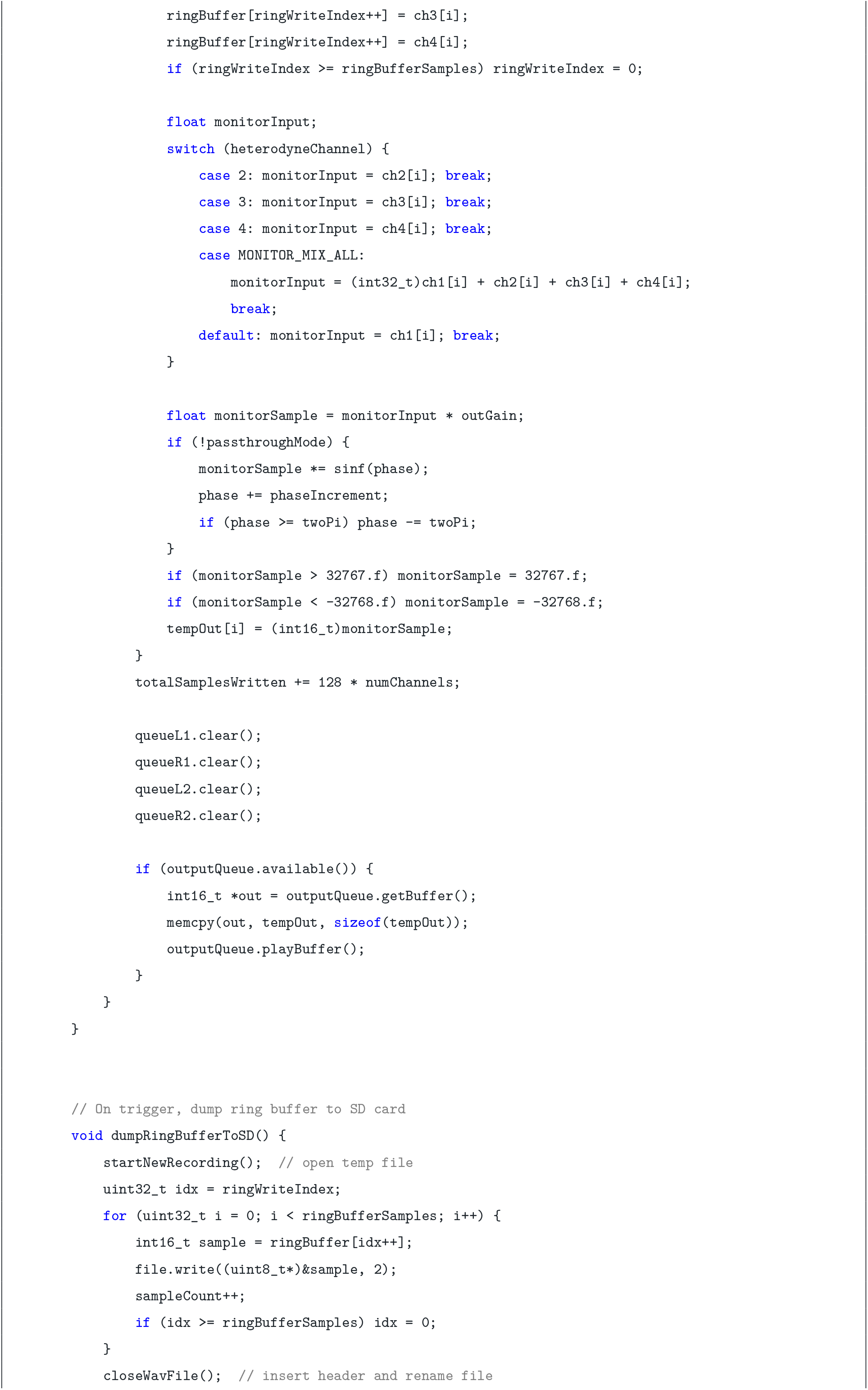
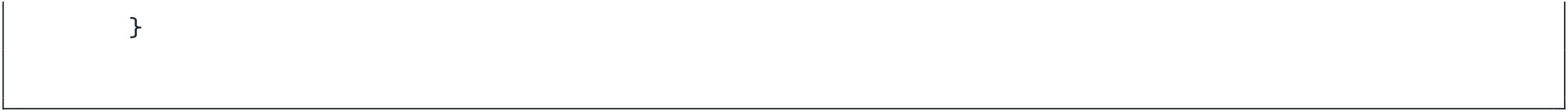
Ring Buffer Implementation

## D REAL-TIME MONITORING IMPLEMENTATION CODE

This appendix provides the key firmware implementation of the selectable real-time monitoring subsystem described in the main manuscript.

### D.1 Global Variables and Constants

Heterodyne mode uses a phase accumulator to generate a continuous sine-wave carrier. The carrier selector also provides passthrough states, and the input selector includes the combined four-channel mix:

The phase increment is calculated during initialisation based on the carrier frequency and sampling rate:

**Listing 3.**
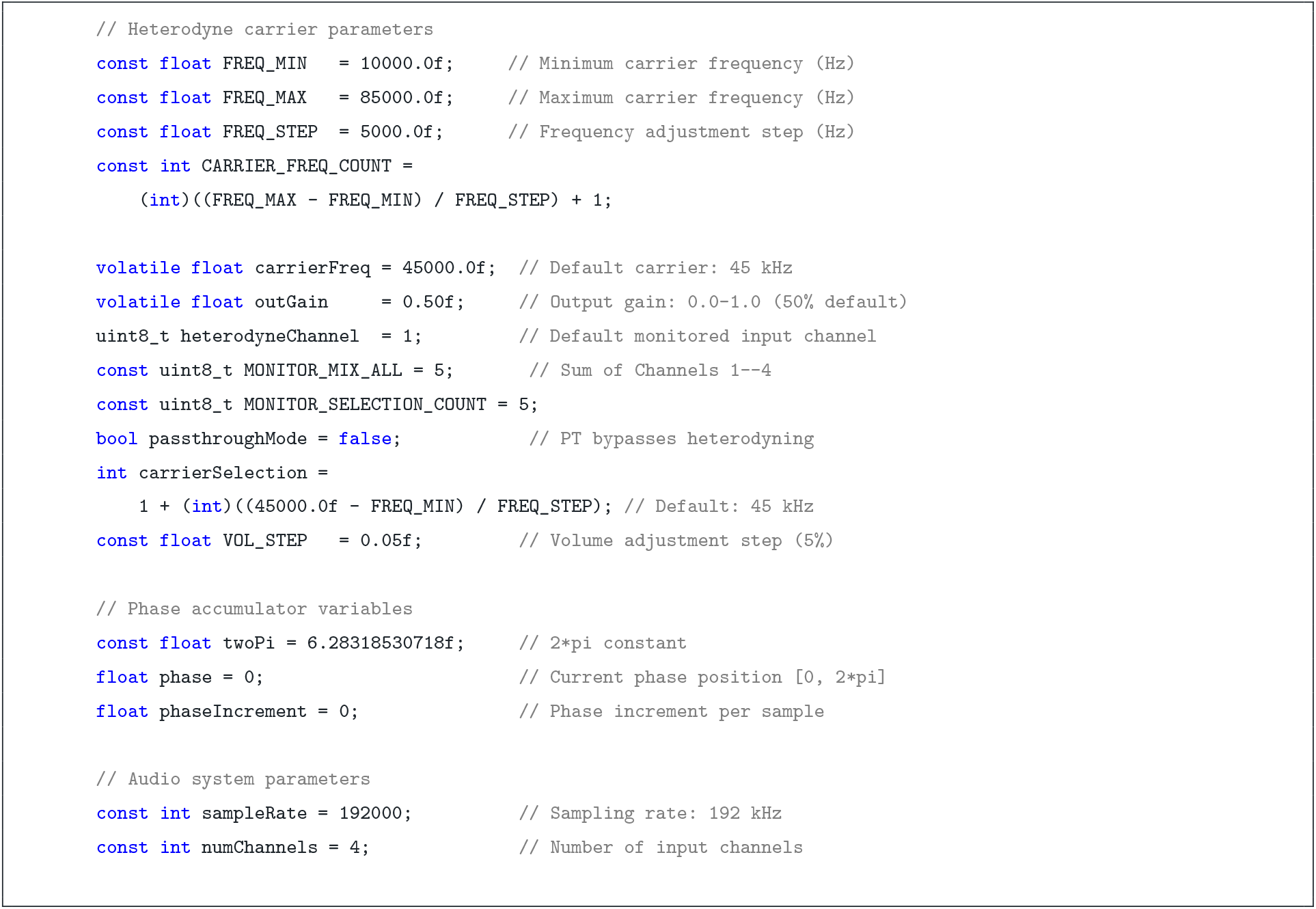
Global variables for selectable real-time monitoring.

**Listing 4.**
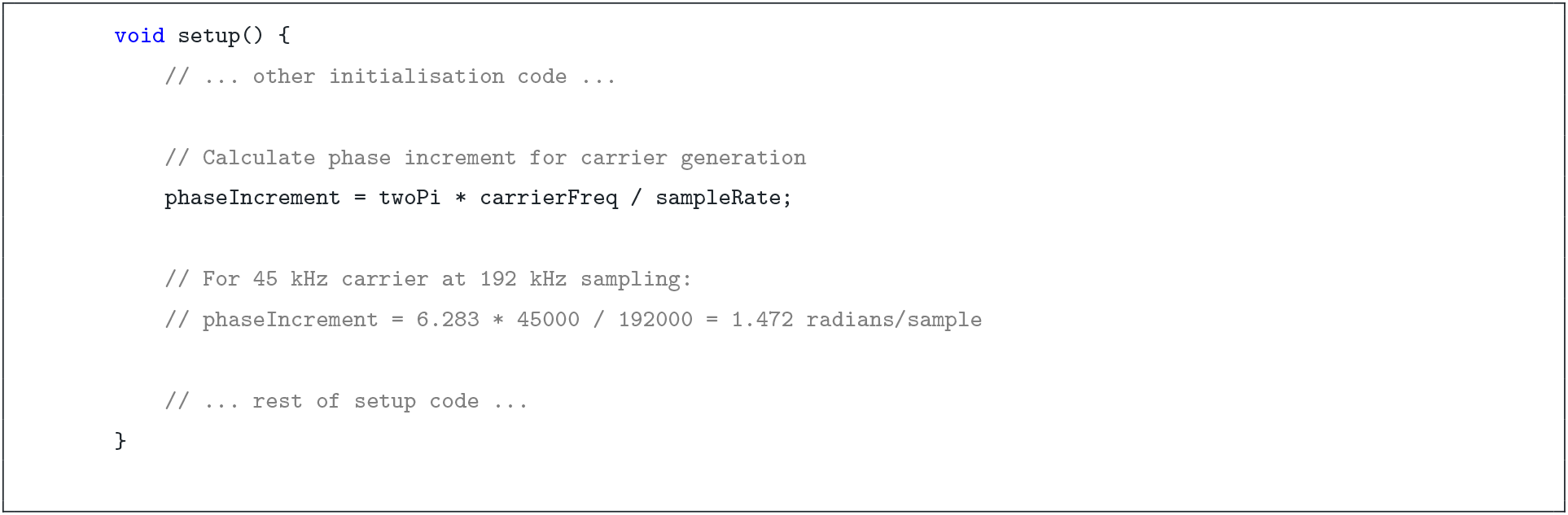
Phase increment initialization in `setup()`.

### D.2 Real-Time Audio Processing

The `readAndProcessAudio()` function fills the PSRAM ring buffer with four raw channels, forms the selected monitor input, and sends the monitoring signal to the DAC. Channel 1 is selected at power-up; the operator can select Channels 1–4 or `MX`, which sums all four inputs. Heterodyne mode multiplies this signal by the carrier, whereas `PT` sends it directly to the output stage.

**Listing 5.**
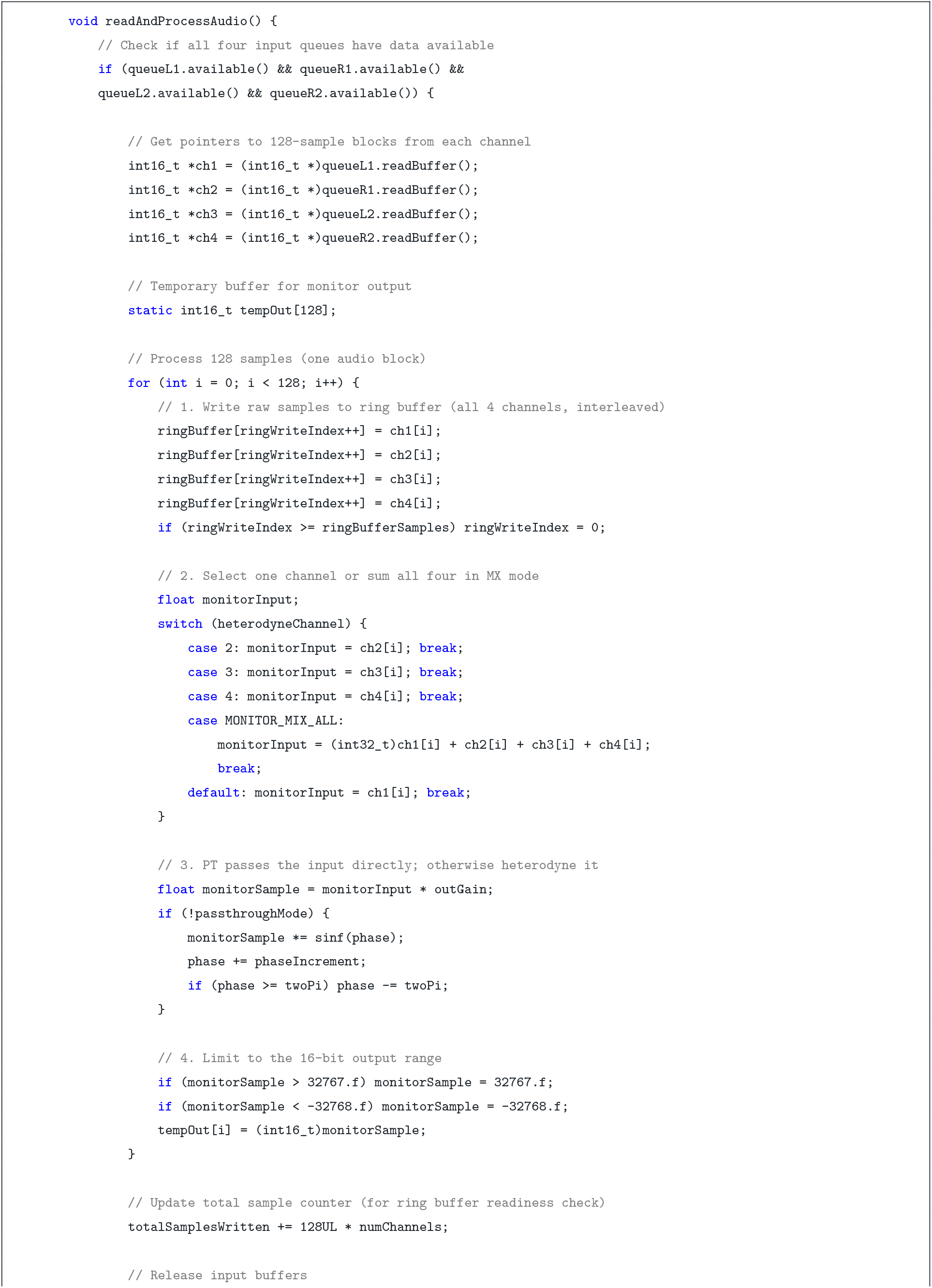
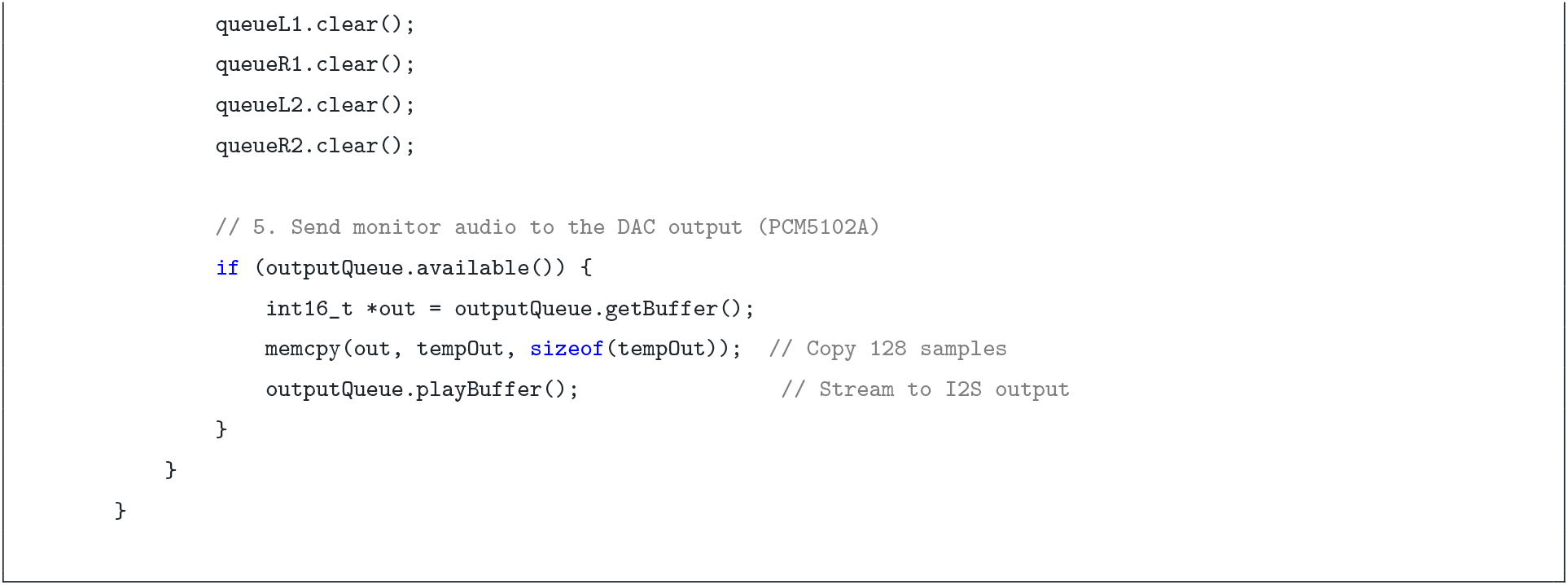
Selectable real-time monitoring with parallel four-channel recording.

### D.3 Monitoring Selection

The rotary-encoder switch cycles among carrier, monitored-input, and output-level controls. The OLED highlights the active control. Carrier selection traverses `PT`, 10–85 kHz, and `PT`; input selection cycles through Channels 1–4 and `MX`:

**Listing 6.**
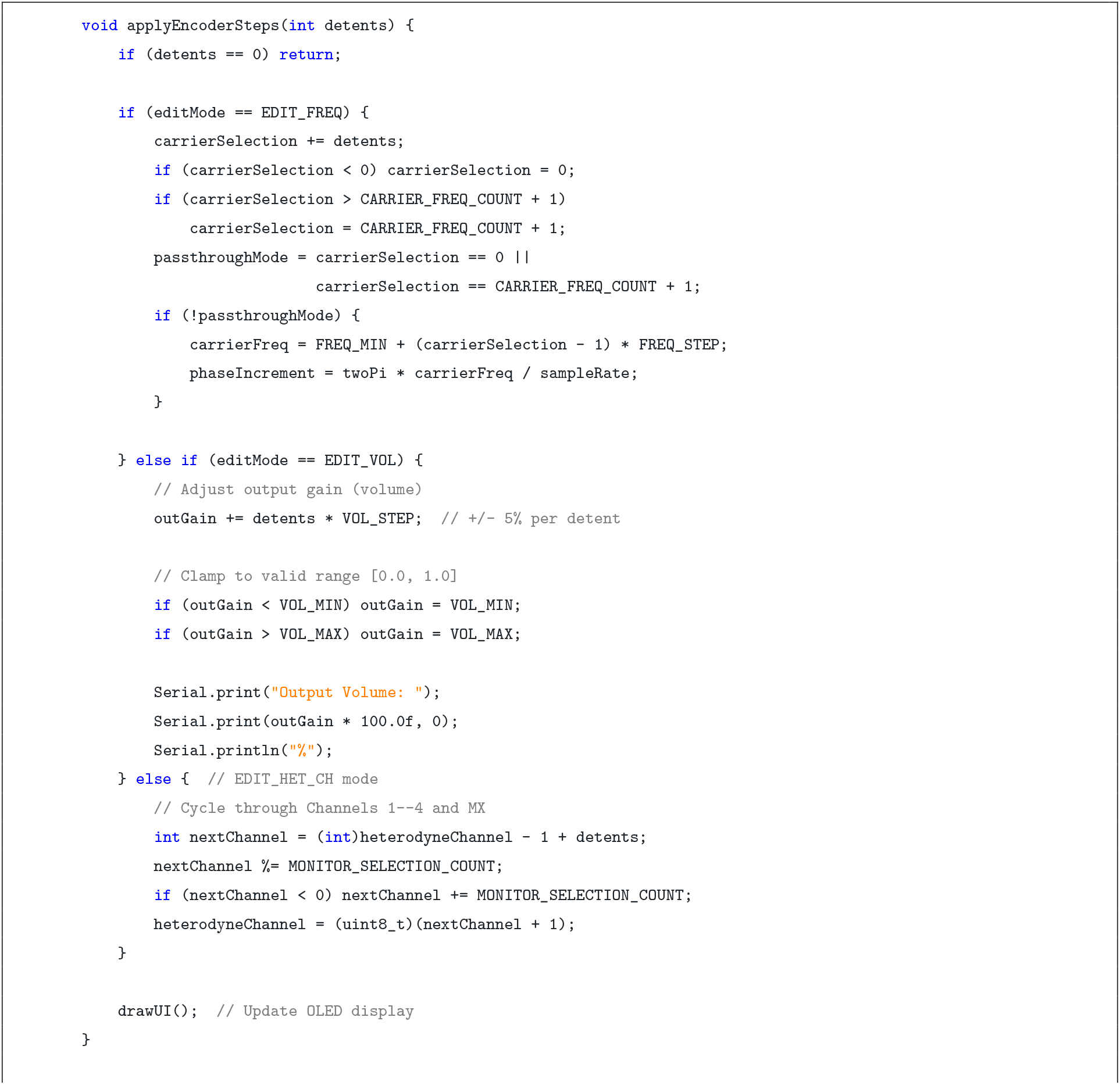
Monitoring-mode and input selection via rotary encoder.

These selections change only the signal sent to the monitoring output. The raw samples from all four microphones continue to be written separately to the ring buffer and saved in every recording.

### D.4 Mathematical Principle

In heterodyne mode, multiplication in the time domain corresponds to convolution in the frequency domain. Given an input signal *x* [*n*] containing a spectral component at frequency *f*_bat_, and a carrier sine wave at frequency *f*_carrier_, the multiplication produces:

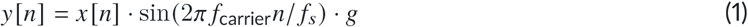

where *f*_*s*_ = 192 kHz is the sampling rate and *g* is the output gain. This operation generates sum and difference frequencies:

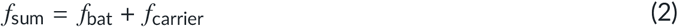

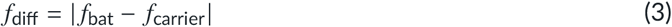

The sum frequency is typically above the Nyquist limit (96 kHz) and is naturally filtered by the DAC reconstruction filter. The difference frequency falls into the audible range, enabling real-time monitoring through standard headphones.

#### Example

A bat echolocation call at 62 kHz mixed with a 60 kHz carrier produces:

- Sum: 62 + 60 = 122 kHz (inaudible, filtered)
- Difference: |62 − 60| = 2 kHz (audible tone)

This 2 kHz tone is heard in real time through the headphones, allowing the researcher to detect bat activity without viewing a spectrogram.

### D.5 Performance Characteristics

- **Computational cost:** Per 128-sample block (0.67 ms at 192 kHz):
  ▪ 128 calls to `sinf()` (hardware-accelerated)
  ▪ 128 floating-point multiplications
  ▪ 128 floating-point to integer conversions
  ▪ Negligible impact on Teensy 4.1 @ 600 MHz
- **Latency:** <1 ms (one audio block buffering delay)
- **Frequency response:** Flat across tunable range (10–85 kHz)
- **Dynamic range:** Limited by 16-bit integer output (∼96 dB theoretical)
- **Harmonic distortion:** Minimal (depends on `sinf()` precision)
- **Interference:** None—operates in parallel with ring buffer recording

## E BILL OF MATERIAL

**Table T2.** Bill of Materials for the Batsy4 Pro Recording System. The listed prices reflect procurement costs in Germany during development, from January to July 2025.

| Item | Description | Count | Unit Cost (€) | Source |
| --- | --- | --- | --- | --- |
| Teensy 4.1 | Microcontroller | 1 | 39.50 | reichelt.de |
| WM8782 | I2S ADC | 2 | 14.90 | [Audio-phonics.fr](https://audio-phonics.fr) |
| Knowles SPU0410LR5H-QB | MEMS Mics | 4 | 0.50 | Digikey.de |
| APS6404L-3SQR PSRAM - 8MB - | External Memory | 2 | 2.50 | tinytronics.nl |
| PC5102A | I2S DAC | 1 | 7.99 | Amazon.de |
| MP1584EN | DC-DC 3A Buck Converter | 1 | 2.00 | Amazon.de |
| SSD1306 | OLED Display | 1 | 5.25 | [AZDelivery.de](https://azdelivery.de) |
| Push Buttons | User Input | 2 | 0.50 | Amazon.de |
| Rotary Encoder | User Input | 1 | 1.50 | Amazon.de |
| Circuit Switch | Power Supply ON/OFF | 1 | 0.25 | Amazon.de |
| Perforated PCB | System Base | 3 | 1.50 | Amazon.de |
| Hexnut Screws | PCB Stacking | Assorted | 5.00 | Amazon.de |
| Wiring | 18/22 gauge coloured wires | Assorted | 5.00 | Amazon.de |
| 6-Pin Connector | Mic power and signal connection | 1 | 14.99 | Amazon.de |
| CR2032 | RTC Battery | 1 | 1.50 | Generic |
| **Total** |  |  | **125.28** |  |

**Table T3.**
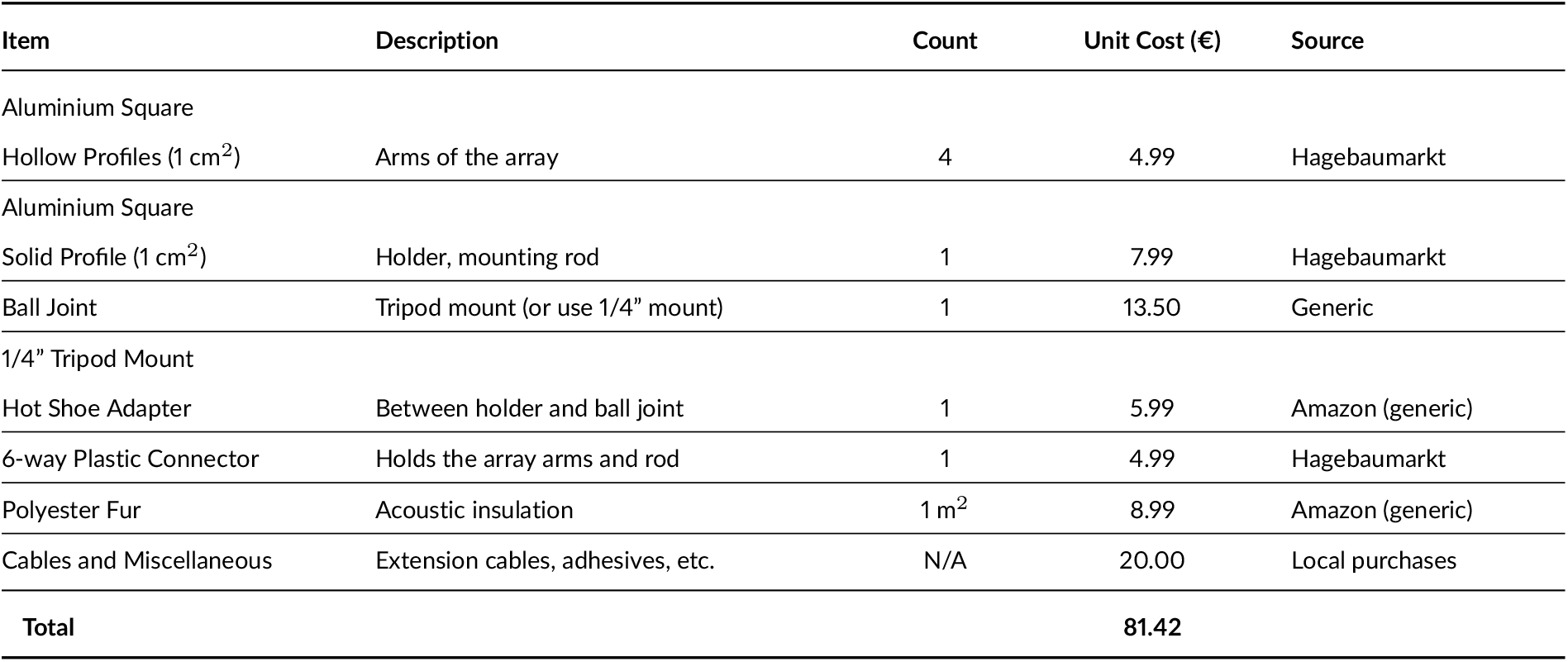
Bill of Materials for the Field Array. The material list is only suggestive; equivalent components may be used depending on array design and local availability.

## F FIELD DATA - CALL DURATION STATS

**Figure S1.**
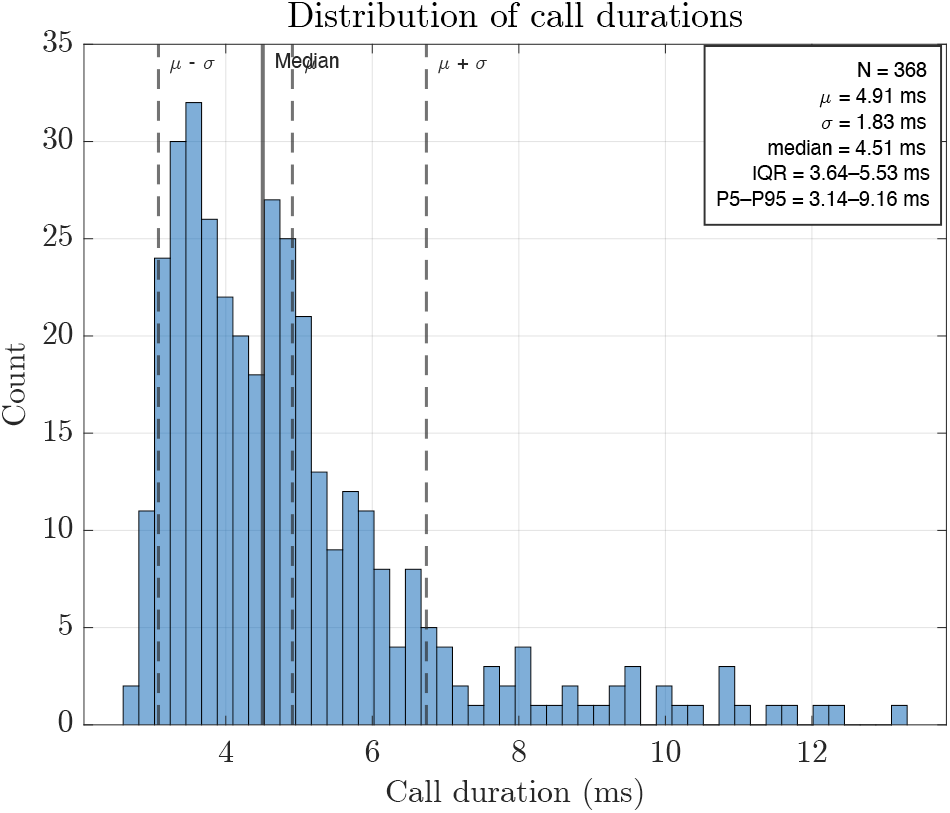
Distribution of echolocation call durations extracted from validated detections (*N* = 368 calls). Call duration was defined using an energy-based percentile criterion: for each call waveform segment, the cumulative sum of squared amplitude was normalised, smoothed, and the call duration was taken as the interval between the time points at which the cumulative energy crossed the lower and upper fractional thresholds (5–95% of total energy). The histogram shows a unimodal distribution centred on short-duration frequency-modulated calls. The vertical solid line indicates the median call duration (4.51 ms), while the shaded region denotes the interquartile range (IQR: 3.64– 5.53 ms). Dashed lines mark the 5th and 95th percentiles (3.14–9.16 ms), illustrating the limited spread and absence of long-duration outliers.

## G GEOMETRY-TO-HUB CONVERSION

**Figure S2.**
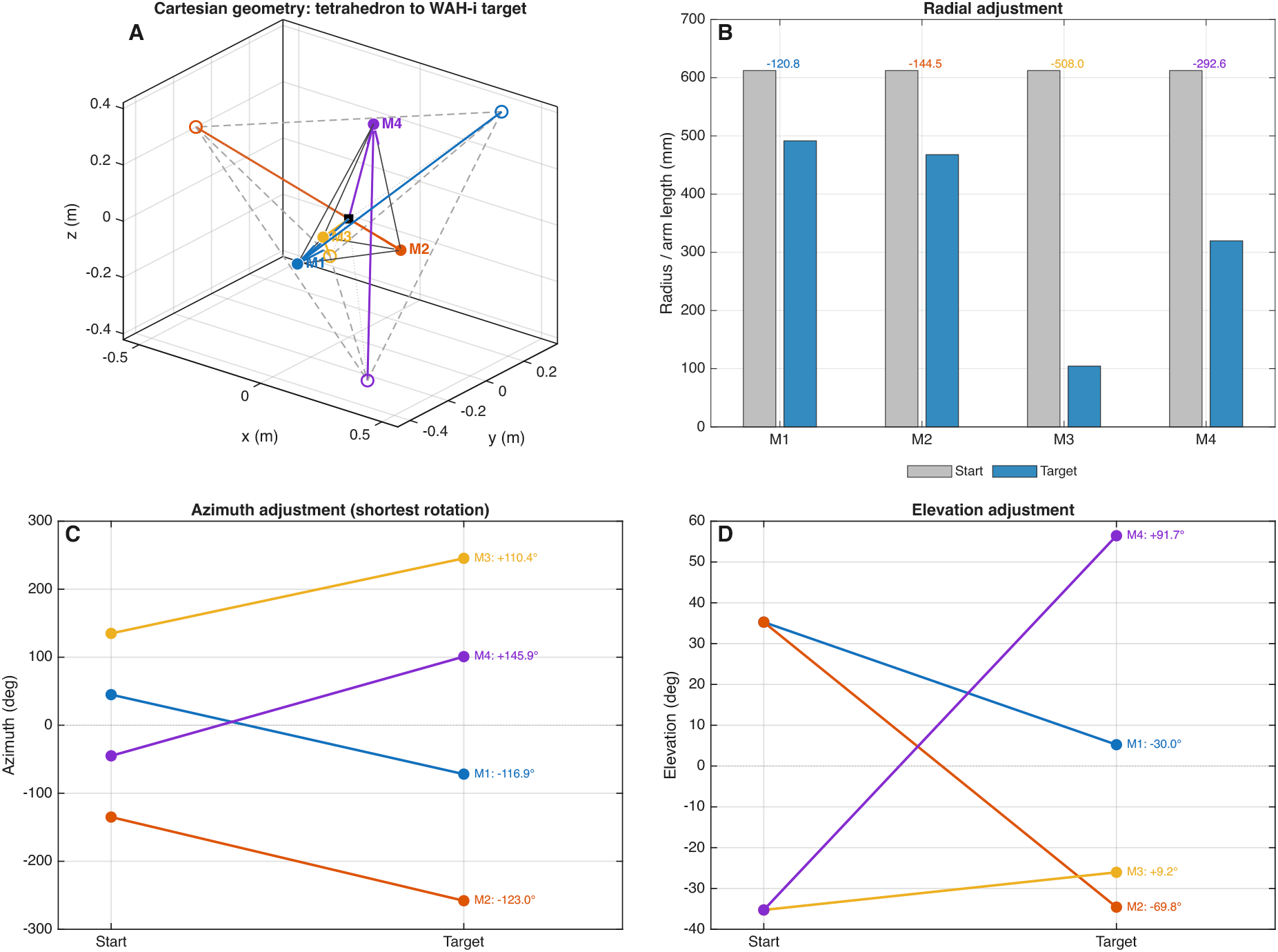
Conversion of an example WAH-*i* Cartesian output into settings for a reconfigurable four-arm hub. (A) Regular tetrahedral starting geometry (open circles and dashed edges; 1 m edge length) and the optimised target geometry (filled circles and solid edges), with arrows indicating microphone displacement. (B) Starting and target radial arm lengths; annotations give the signed length changes in millimetres. (C) Starting and target azimuths, expressed using the shortest signed rotations. (D) Starting and target elevations. Coordinates share a common hub origin, with +*x* defining zero azimuth, +*y* positive azimuth, and +*z* positive elevation; microphone identity is preserved by CSV row order.

## Footnotes

1 `[OSHW] DE000165 | Certified open source hardware` | oshwa.org

